# Reprogramming a thiosulfate bacterial two-component system into a perchlorate biosensor

**DOI:** 10.64898/2026.11.13.766565

**Authors:** Samuel Lim, James J. McCann, Piyush Kumar, Zhujin Xia, Paul Molinaro, William R. Henson, Ronald L. Koder, Scott Banta

## Abstract

Engineering bacterial two-component systems (TCSs) to sense new molecules would enable programmable whole-cell biosensors, but retargeting membrane-bound sensor histidine kinases is challenging because ligand recognition and signal transduction are tightly coupled. Here, we reprogram the thiosulfate-responsive TCS from *Shewanella halifaxensis* (ThsSR) to detect perchlorate, for which no naturally occurring TCS sensor has been reported. Using a dual growth selection, we simultaneously selected for perchlorate-dependent signaling and against native thiosulfate responsiveness. This strategy first converted wild-type ThsS into a perchlorate-responsive generalist and then produced specialized variants with improved sensitivity, greatly reduced thiosulfate response, and discrimination against related oxyanions. We next applied the physics-informed reinforcement learning design method protCAD to functional intermediates identified by directed evolution. protCAD-selected substitutions further enhanced sensitivity or signaling output in specialized sensors, even in backgrounds that were refractory to additional random mutagenesis and gene shuffling. As a final demonstration, we used the engineered TCS sensor in a whole cell biosensor to detect perchlorate in unsterile environmental samples. Together, these results define a productive division of labor between experiment and computation: directed evolution discovers signaling-competent, specificity-altered sequence space, while computation explores that evolved space more completely to refine performance. This evolution-computational strategy provides a framework for reprogramming TCS biosensors toward non-native analytes.

**Significance:** Protein design and directed evolution are often framed as alternative paths to engineered function. We show instead that they are most powerful when assigned different tasks. Directed evolution transformed a membrane-bound thiosulfate sensor into a perchlorate-responsive generalist, and subsequent counterselection removed the native thiosulfate response. Structure-based computational design was then applied to these evolved, signaling-competent intermediates and prospectively identified mutations that further improved whole-cell performance. The result is a division of labor: evolution finds functional sensor architectures and enforces specificity in cells, whereas computation searches the evolved sequence space more deeply to tune sensitivity, selectivity, and output. This evolution-guided design strategy establishes a route to reprogram bacterial two-component systems toward non-native analytes, including targets with no known natural sensor.

## Introduction

Bacteria have evolved the ability to sense and respond to a wide range of environmental stimuli. Signals detectable by bacteria include biochemical cues such as small molecules and polymers, as well as physical conditions including pH, temperature and light (1). Synthetic biologists have been keen to reprogram such capabilities to engineer genetically encoded biosensors. These microbial biosensors have proven useful for diverse applications including environmental monitoring (2), metabolic production (3) and biomedicine (4). Bacterial biosensors typically rely on either transcription factors or riboswitches to generate output signals upon detection of the target stimulus (1). Transcription factors regulate the reporter gene expression by selectively occupying promoter regions upon allosteric ligand binding (5), while riboswitches function by altering mRNA structure in response to the ligand binding to an aptamer (6). However, both systems can only respond to internal cytoplasmic conditions, limiting the possibilities of biosensor design as many external signals cannot penetrate cell membranes (7).

Bacterial two-component systems (TCSs) represent a promising alternative, as they constitute a signal transduction pathway capable of detecting extracellular stimuli and subsequently generating intracellular responses (7). A typical TCS consists of three different components including a sensor histidine kinase (SHK), a response regulator (RR) and a downstream output promoter (8). In these systems, the SHK functions as a membrane-bound detector module. Ligand binding to its N-terminal extracellular (or periplasmic) domain triggers autophosphorylation in the cytoplasmic C-terminal domain. The phosphoryl group is then transferred to the RR, activating it to modulate the expression of genes controlled by the output promoter (9). In general, SHKs tend to be highly specific towards their cognate RRs, ensuring minimal crosstalk with other pathways within a crowded cellular environment (10). Because of this, while RRs are conserved, SHKs are modular and highly variable in sequence (11). Rewiring diverse types of TCSs to desired output genes resulted in microbial cells that can be patterned onto material surface in a light-responsive manner (12), or those capable of sensing inflammatory biomarkers upon administration into the mouse gut (13).

A major bottleneck in exploiting TCSs for biosensor development is the paucity of well-characterized systems: although over 160,000 genes from more than 2,000 prokaryotic genomes have been annotated as encoding TCS proteins (14) only small fraction of these TCSs, on the order of few hundred, have experimentally identified input signals (7). Target screening of these “orphan TCSs” is time-consuming and often limited by genetic intractability of their host organisms. Therefore, to expand the applicability of TCS-based biosensors it is desirable to develop methods to engineer the detection specificity of known TCSs towards new target molecules.

Reprogramming a TCS to recognize a non-native signal remains challenging because SHK input domains are poorly conserved, structurally undercharacterized in both signaling states, and difficult to relate mechanistically to output from the cytoplasmic kinase domain. The modular architecture of TCSs has enabled one successful strategy: replacing an SHK input domain with that from an orthogonal sensor. These domain-swap chimeras can rewire endogenous pathways in model organisms such as *E. coli*, allowing native TCS circuitry to respond to alternative signals without heterologous expression of an entire non-native pathway (15-19). However, this approach is limited to exchanging specificities among naturally occurring sensors. It does not solve the more general problem of directing an existing TCS to recognize a genuinely new molecular target.

Directed evolution provides a direct route to new protein function by iteratively coupling genetic diversification to phenotypic selection (20). This is especially valuable for protein–ligand recognition, where productive substitutions often occur outside obvious contact residues and are difficult to predict *a priori* (21-23). Consequently, directed evolution has been used to retarget diverse enzymes (24-26) and transcription factors (27, 28) toward non-native substrates and ligands. In complex multiconformational signaling proteins such as SHKs, directed evolution can also provide a different kind of information: it can empirically define which regions of the sensor tolerate mutation while preserving signal transmission.

Computational protein design offers a complementary way to exploit that experimentally defined mutational space. Recent advances in computational design have also revolutionized the field of protein engineering (29-31). Structure-prediction methods such as AlphaFold3 (32), RoseTTAFold (33) and ESMFold (34) together with generative and sequence-design models including RFdiffusionAA (33, 35), ProGEN3 (36), LigandMPNN (37), and ESM3 have enabled the design of proteins that bind new small-molecule targets (38-40). Although computational methods are often used to nominate mutation sites before experimental screening (41-43), our strategy reverses this logic: directed evolution first identifies the sensor regions compatible with functional signaling, and computation is then used to explore that permissive sequence space more completely than would be practical by experimental library screening alone. This evolution-design cycle enables broader coverage of viable sequence space while retaining the functional constraints imposed by the native TCS architecture.

Here, we reprogram the ligand specificity of the thiosulfate-sensing TCS ThsSR from *Shewanella halifaxensis* (13) to detect perchlorate, a target for which no naturally occurring TCS sensor has been reported. When initial attempts at computational redesign failed, we developed a dual growth-based selection that couples positive selection for perchlorate responsiveness with negative selection against native thiosulfate responsiveness, enabling simultaneous gain of the new input and loss of the original one. After directed evolution identified functional ThsS variants and defined regions permissive to mutation, we applied our reinforcement-learning design algorithm, protCAD (44, 45), to more comprehensively explore this tolerated sequence space. This combined strategy yielded point mutations that further increased perchlorate sensitivity and signaling output. After demonstrating perchlorate and signaling output in controlled settings, we engineered a whole cell biosensor using our best performing perchlorate sensing system and demonstrated perchlorate detection in unsterile environmental samples with spiked perchlorate. This work establishes a generalizable evolution/computational strategy for converting natural TCSs into highly specific microbial biosensors for non-natural inputs.

## Results and Discussion

### Initial attempts at computational design

We started by surveying structurally characterized TCS SHKs for ligand-binding domains that recognize oxyanions similar to perchlorate, with the goal of minimizing the sequence changes required for retargeting. Our first candidate was NarQ, a nitrate-responsive TCS SHK that has been extensively structurally characterized (9). However, NarQ proved unsuitable: modeling indicated that replacing the planar nitrate anion with tetrahedral perchlorate would require substantial backbone rearrangement to accommodate the larger, nonplanar ligand. We therefore turned to the thiosulfate-responsive ThsSR system from *Shewanella halifaxensis* (13), because its cognate ligand more closely resembles perchlorate in size and tetrahedral geometry. Although this ThsSR has been functionally characterized, no experimentally determined apo- or holo-state structures are available either for ThsS or its periplasmic sensor domain. Nevertheless, AlphaFold 3 modeling of the ThsS–thiosulfate complex suggested that the binding pocket could be reprogrammed for perchlorate recognition without extensive backbone remodeling. The predicted structure of the ThsS–thiosulfate complex is shown in Figure 1a.

**Figure 1.**
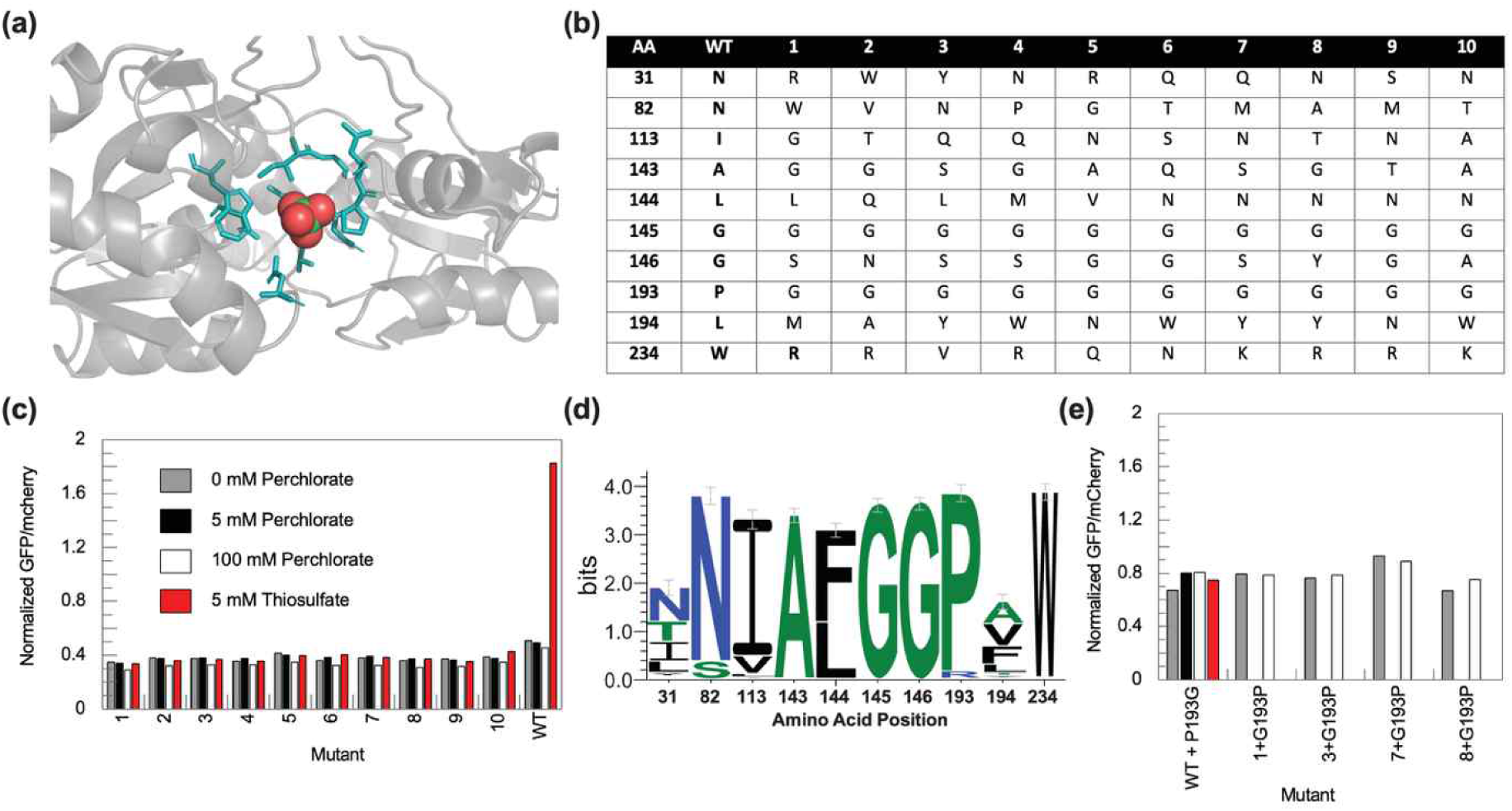
Computational redesign and functional screening of ThsS variants designed for perchlorate recognition. **(a)** AlphaFold 3-predicted structure of the *Shewanella halifaxensis* ThsS periplasmic sensor domain bound to thiosulfate. The protein backbone is shown in gray, the ten first- and second-shell residues selected for redesign are shown as cyan sticks, and the modeled thiosulfate ligand is shown as spheres. **(b)** Amino acid identities at the ten redesigned positions in wild-type (WT) ThsS and the ten highest-scoring sequences generated by protCAD. **(c)** Functional screening of WT ThsS and designs 1–10 in the *Escherichia coli* ThsSR reporter platform. ThsR-dependent sfGFP fluorescence was normalized to constitutive mCherry fluorescence following growth with no added ligand (grey), 5 mM perchlorate (black), 100 mM perchlorate (white), or 5 mM thiosulfate (red). WT ThsS exhibited thiosulfate-dependent activation, whereas none of the designed variants exhibited detectable ligand-dependent activation by either thiosulfate or perchlorate. The data shown are from a single preliminary experiment. **(d)** Sequence logo for the ten redesigned positions, generated from an alignment of 250 two-component-system sensor histidine kinases with diverse ligand specificities. Total stack height represents positional information content in bits, and the relative height of each letter within a stack represents amino acid frequency. **(e)** Functional analysis of residue Pro193. The P193G substitution was introduced into WT ThsS and assayed under all four conditions shown in panel C. G193P revertants of designs 1, 3, 7, and 8 were assayed without added ligand and with 100 mM perchlorate. WT ThsS P193G did not retain the thiosulfate-dependent response of WT ThsS, and restoring Pro193 in the selected designs did not produce perchlorate-dependent activation. The data shown are from a single preliminary experiment.

Using this model, we selected ten first- and second-sphere residues surrounding the modeled thiosulfate ligand as targets for computational optimization using our physics-informed reinforcement learning design method protCAD (Figure S1a) (45). We chose the 10 best scoring sequences (Figure 1b) and screened each for ligand retention using 25 ns MD simulations. All ten retained the ligand, although in some cases the ligand moved several angstroms from their initial postion during the simulations (Figure S1b). We introduced these sequences into an *E. coli* sensing platform comprised of the plasmids pKD182 and pKD184 created by Tabor and coworkers (Figure S2) (13). pKD182 expresses the transmembrane SHK sensor protein ThsS under the control of an IPTG-inducible tac promoter while pKD184 contains the superfolder green fluorescent protein (sfGFP) under the control of ThsR-activated promoter P_phsA_ as well as constitutively expressing (i) RR element ThsR, (ii) mcherry, (iii) the antibiotic resistance proteins tetR and (iv) cmR. Ligand binding to the extracellular domain of ThsS triggers phosphorylation of ThsR and the activated ThsR binds to the PphsA promoter and induces sfGFP expression, enabling convenient expression level-normalized measurement of TCS response using the GFP-to-mcherry fluorescence ratio.

Figure 1c shows the normalized sfGFP/mCherry output of wild type (WT) ThsS and the ten designed variants in the absence of added ligand and following treatment with 5 mM thiosulfate or 5 or 100 mM perchlorate. WT ThsS exhibited a strong thiosulfate-dependent response but no detectable response to perchlorate. In contrast, none of the designed variants showed ligand-dependent activation by either thiosulfate or perchlorate. The absence of reporter induction does not distinguish between loss of ligand recognition, impaired protein expression or folding, or disruption of signal transmission through the receptor. Given the general degree of success of the protCAD algorithm, it seems likely that at least some of the variants may bind perchlorate but do not transmit a signal. We therefore examined sequence conservation at the redesigned positions across 250 homologous TCS SHKs (Figure 1d). Several of these positions were strongly conserved, most notably Pro193, which protCAD replaced with glycine in all ten perchlorate-targeted designs. Introducing the P193G substitution into otherwise WT ThsS abolished its thiosulfate-dependent response (Figure 1e), demonstrating that Pro193 is required for WT sensor function under these conditions. However, restoring Pro193 in a subset of the designed variants did not confer perchlorate-dependent activation, indicating that the inactivity of these variants cannot be attributed to the P193G substitution alone. These results suggest that successful ligand retargeting will require preserving residues that support receptor structure and signaling while optimizing interactions within the ligand-binding pocket. However, due to the lack of structural information on this TCS the identities of these residues are unknown.

### Growth-based selection and directed evolution experiments

Evolutionary approaches enable a broader survey of the sequence space accessible to optimization in the full signal transduction apparatus, and both serve to scan for improved function and determine which residues are amenable to alteration. Thus, we explored directed evolution to shift the ligand specificity of ThsSR system from thiosulfate to perchlorate. Importantly, our goal was not only to engineer ThsSR to recognize perchlorate but also to suppress its native response to thiosulfate. To achieve this, we employed a dual-selection system, during which positive selection cycles enrich cells expressing perchlorate-responsive TCS variants and negative selection cycles exclude those retaining response to thiosulfate or those showing elevated basal activity (Figure 2a). To do this we replaced pKD184 with pKD184-dual (Figure S2), which included genes encoding kanamycin resistance (*kanR*) and sucrose sensitivity (*sacB*) to the sfGFP under the control of ThsR-activated promoter P_phsA_. When ligand binding to the extracellular domain of ThsS triggers phosphorylation of ThsR, the activated ThsR binds to the PphsA promoter and induces *kanR* expression, conferring kanamycin resistance and enabling positive selection in the presence of the target ligand perchlorate. In contrast, the expression of sacB gene under the presence of sucrose is lethal, as levansucrase converts sucrose into the toxic polymer levan (46). Therefore, sucrose addition to the culture media enables negative selection against the variants that constitutively activate the P_phsA_ promoter, while the simultaneous addition of sucrose and native ligand (thiosulfate) eliminates mutants that remain active towards thiosulfate. We note that in this design *sacB* would be expressed during the positive selection as well, but there will be no effect when sucrose is absent in the media.

**Figure 2.**
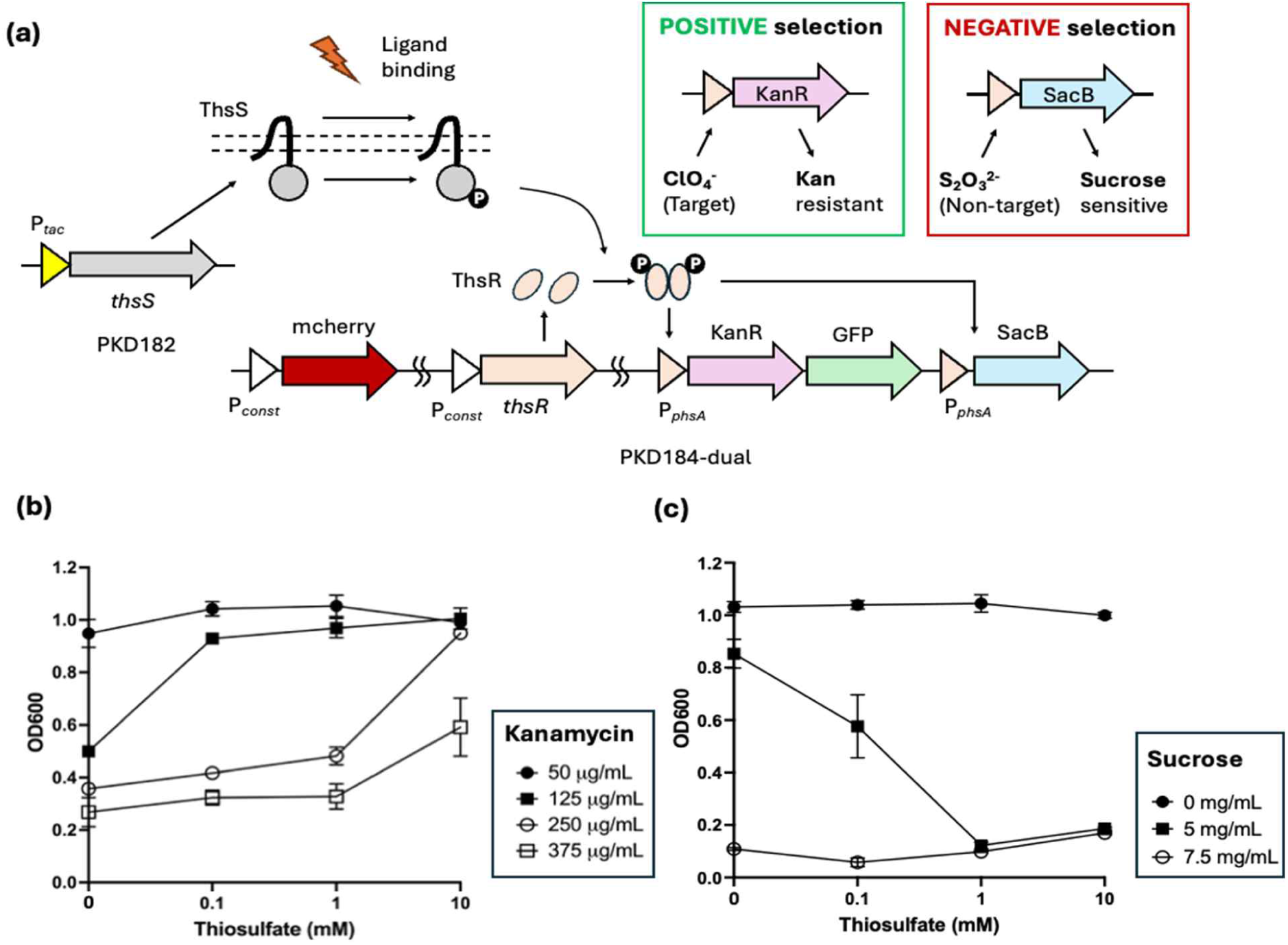
Design of the growth-based dual selection system for the directed evolution of TCS biosensor. **(a)** Plasmid PKD182 expresses ThsS sensor protein under the IPTG-inducible tac promoter, and plasmid PKD184-dual contains constitutively expresses mCherry and the ThsR response regulator gene as well as the KanR, GFP and sacB genes under the control of TCS ligand-activated P_phsA_ promoter. Activation of the KanR gene upon addition of perchlorate allows for positive selection under kanamycin, whereas expression of sacB upon addition of thiosulfate leads to negative selection under sucrose. **(b)** Growth of cells containing dual-selection system as a function of the concentration of kanamycin and thiosulfate. OD measurements were taken after 12 hours of incubation. Growth of cells were promoted upon increasing concentration of thiosulfate at 125 – 375 µg/mL kanamycin. Each data point indicates average from three independent experiments (N=3), and the error bars indicate standard deviation. **(c)** Growth of cells containing dual-selection system under varying amounts of sucrose and thiosulfate concentrations. The OD measurements were taken after 12 hours of incubation. Growth of cells were inhibited upon increasing concentration of thiosulfate at 5 mg/mL. Each data point indicates average from three independent experiments (N=3), and the error bars indicate standard deviation.

To validate the selection system and optimize culture conditions, *E. coli* cells co-transformed with PKD182 and PKD184-dual were grown in media containing varying concentrations of kanamycin or sucrose, and their growth was measured under different thiosulfate concentrations. In an experiment to test the positive selection condition, cell growth under kanamycin selection was enhanced with increasing concentration of thiosulfate from 0.01 mM to 10 mM (Figure 2b, Figure S3a-e). This trend persisted at kanamycin concentrations tested (125 - 375 µg/mL) except for 50 µg/mL, indicating that the system functions robustly under stringent selection conditions. Furthermore, it was demonstrated that the selection could be carried out on solid agar plates containing kanamycin and thiosulfate (Figure S4). When sucrose was added to 5 mg/L, the addition of thiosulfate increasingly suppressed cell growth, demonstrating that the negative selection system effectively excludes variants exhibiting basal level activation of the TCS (Figure 2c, Figure S3f-g). Thus, selection conditions were set at 375 µg/mL kanamycin and 5 mg/mL sucrose for the positive and negative cycles, respectively.

Directed evolution experiments were conducted in three iterative cycles as described in Figure 3a. Each cycle started with construction of the mutagenesis library of PKD182 plasmid, where mutations were introduced into *thsS* gene. For each round different mutational methods were explored including site-saturation mutagenesis, random mutagenesis and gene shuffling, which are described in detail in later sections. Libraries were transformed into *E. coli* cells previously transformed with the PKD184-dual plasmid, cultured the transformed cells in liquid media, and subsequently selected them on kanamycin-agar plates containing different concentrations of perchlorate as the positive selection cycle. Selected colonies were grown from agar plates with lowest perchlorate concentration from each cycle to proceed to the next step. To avoid losing beneficial mutations prematurely through stringent selection, negative selection was not applied in the initial iteration cycle. For the second and third iterations, colonies obtained from the positive selection were collected and incubated in the presence of sucrose to remove the pseudo-positive variants with high non-specific responses, and subsequently plated on agar plates containing no sucrose or kanamycin. Colonies obtained from either positive or negative selection were individually screened for their ability to recognize perchlorate. Successful variants were then sequenced to identify the mutations, and the corresponding clones were subsequently used as templates for the next round of mutagenesis. Details regarding each selection cycle are outlined in Table S1.

**Figure 3.**
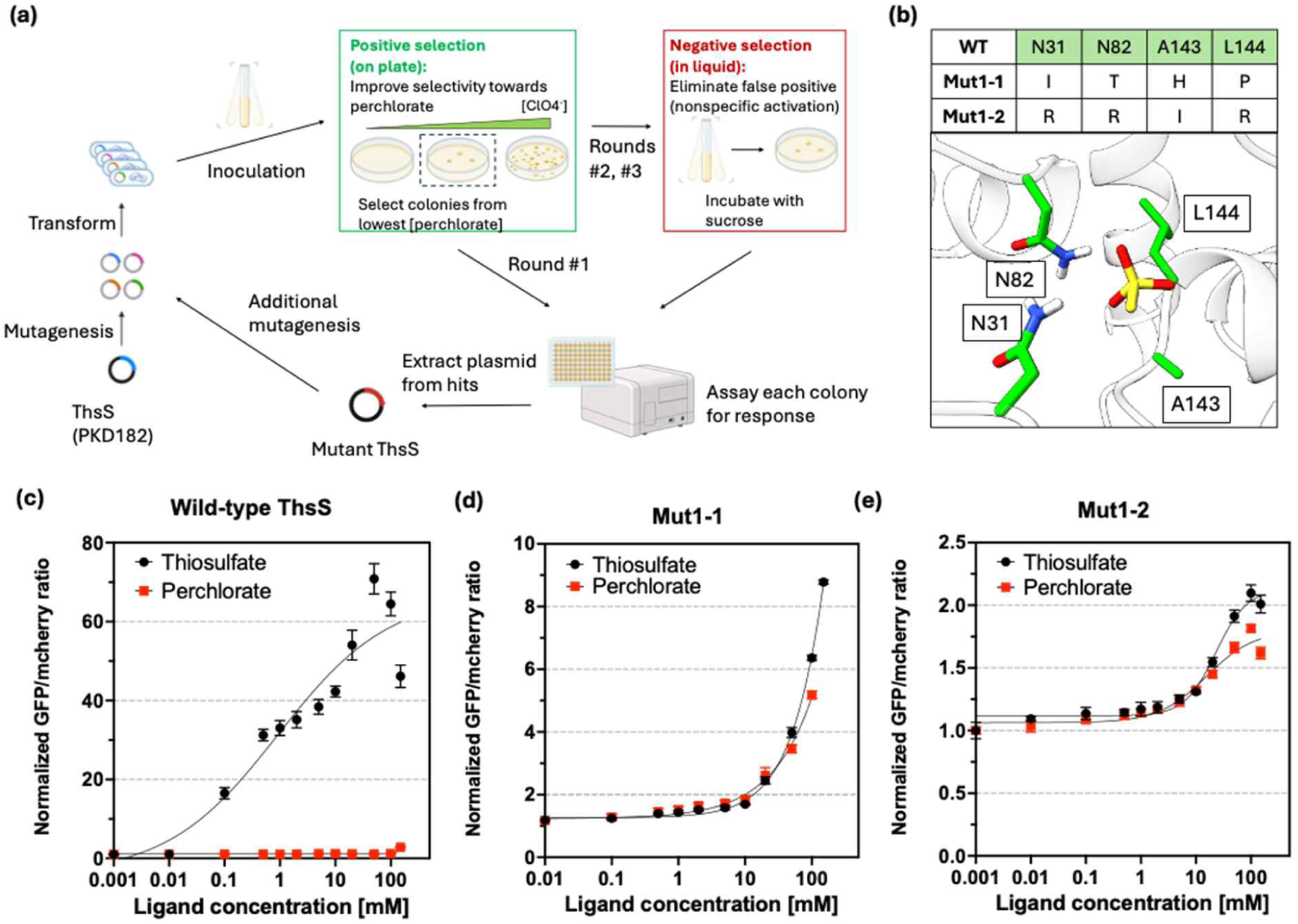
Site-saturation mutagenesis generates promiscuous variants of ThsS. **(a)** Experimental steps for the directed evolution of the ThsS TCS towards detection of perchlorate. For each iteration, mutant library was constructed through *in vitro* mutagenesis and subsequently transformed into *E. coli* cells. For the first round of mutation, only the positive selection was performed to avoid premature loss of potentially beneficial mutations. For the second and third rounds, negative selections were applied to eliminate the false positives and/or the variants having high basal activities. **(b)** Four residues N31, N82, A143 and L144, which were predicted to be within < 4 Å distance from the potential thiosulfate binding site of the wild-type ThsS, were selected as the initial mutagenesis targets to construct site-saturation library. Growth-based positive selection isolated two variants, Mut1-1 and 1-2, as the positive hits with perchlorate responses. Dose-response curves for the **(c)** WT ThsS, **(d)** Mut1-1 and **(e)** Mut1-2 are shown. In contrast to the WT, which is highly specific towards thiosulfate, both mutants display promiscuity, with similar responses towards thiosulfate and perchlorate. Due to the large differences in the dynamic range of each variant, the y-axes of each graph is depicted in different scales. GFP/mCherry response signals were normalized with respect to the basal response of each variant without ligand addition. Each data point indicates average from three independent experiments (N=3), and the error bars indicate standard deviation.

### Cycle 1: Generating a ligand-promiscuous ThsS intermediate by site-saturation mutagenesis

Directed evolution campaigns to alter molecular specificity often benefit from first targeting residues within the binding pocket and then exploring more distant positions in subsequent rounds (22). We therefore began with a semirational site-saturation library focused on residues predicted to contact the ligand directly. Of the ten positions considered in the initial computational design, six—N31, N82, A143, L144, G145, and G146—were located within 4 Å of the modeled thiosulfate ligand in WT ThsS (Figure 3b). We selected N31, N82, A143, and L144 for randomization while retaining G145 and G146 because these adjacent glycine residues may contribute to the local backbone geometry and flexibility of the binding pocket.

An NNK library targeting these four positions was transformed into *E. coli* harboring pKD184-dual. The resulting library contained approximately 10^6^ independent transformants, providing nominally greater than sixfold sampling of the 20^4^=160,000 possible amino acid combinations. Positive selection on agar containing 375 µg/mL kanamycin and 1, 10, or 100 mM perchlorate produced colonies at 10 and 100 mM perchlorate but not at 1 mM. Individual colonies recovered from the 10 mM perchlorate plate were screened using the ratiometric sfGFP/mCherry reporter assay. Two variants exhibited clear perchlorate-dependent activation: Mut1-1 (N31I/N82T/A143H/L144P) and Mut1-2 (N31R/N82R/A143I/L144R).

Dose-response characterization of WT ThsS and the two variants is shown in Figures 3c–e. WT ThsS responds to thiosulfate but showed no detectable activation by perchlorate at concentrations up to 100 mM. In contrast, both Mut1-1 and Mut1-2 exhibit increasing reporter activation between 10 and 100 mM perchlorate while retaining responsiveness to thiosulfate. Thus, the first round of evolution generated ligand-promiscuous sensors rather than fully switching ligand specificity. The high perchlorate concentrations required for activation indicate that both variants initially had low sensitivity to the new ligand. Promiscuous variants with even modest activity can provide productive intermediates from which increased sensitivity and specificity subsequently evolve (47, 48). Thus, Mut1-1 was selected as the parent for further evolution because it exhibited lower basal activity and exhibited a larger inducible range than Mut1-2. Subsequent rounds were therefore designed to increase perchlorate sensitivity while reducing the residual response to thiosulfate. Because perchlorate toxicity prevents reliable characterization at higher concentrations, the Mut1-1 dose-response analysis was limited to concentrations of 100 mM or below.

### Cycle 2: Increasing perchlorate sensitivity and functional selectivity through random mutagenesis

For the second evolutionary cycle, the entire periplasmic sensor domain coding sequence of Mut1-1 was subjected to error-prone PCR, allowing beneficial mutations to arise both within and beyond the modeled ligand-binding pocket. The resulting library was positively selected on agar containing 375 µg/mL kanamycin and 1, 10, or 100 mM perchlorate. Whereas the first-round library produced colonies only at 10 and 100 mM perchlorate, the second-round library also produced colonies at 1 mM, consistent with enrichment of variants capable of activating the pathway at lower perchlorate concentrations. Colonies from the 1 mM plate were pooled, subjected to sucrose counterselection in liquid culture to eliminate variants with elevated ligand-independent activity, and then replated to isolate individual clones for characterization.

Four second-round variants, Mut2-1 through Mut2-4, were isolated. All four retained the four substitutions present in Mut1-1 and acquired two or three additional substitutions distributed throughout the sensor domain (Figure 4a–b). Each variant exhibited response at lower perchlorate concentrations relative to Mut1-1, with activation observed over the 0.1–10 mM concentration range (Figure 4c). Their basal reporter outputs were also lower than those of Mut1-1 and Mut1-2; the basal output of Mut2-4 approached that of WT ThsS (Figure S4). This reduction is consistent with the intended effect of sucrose counterselection against constitutively active variants.

**Figure 4.**
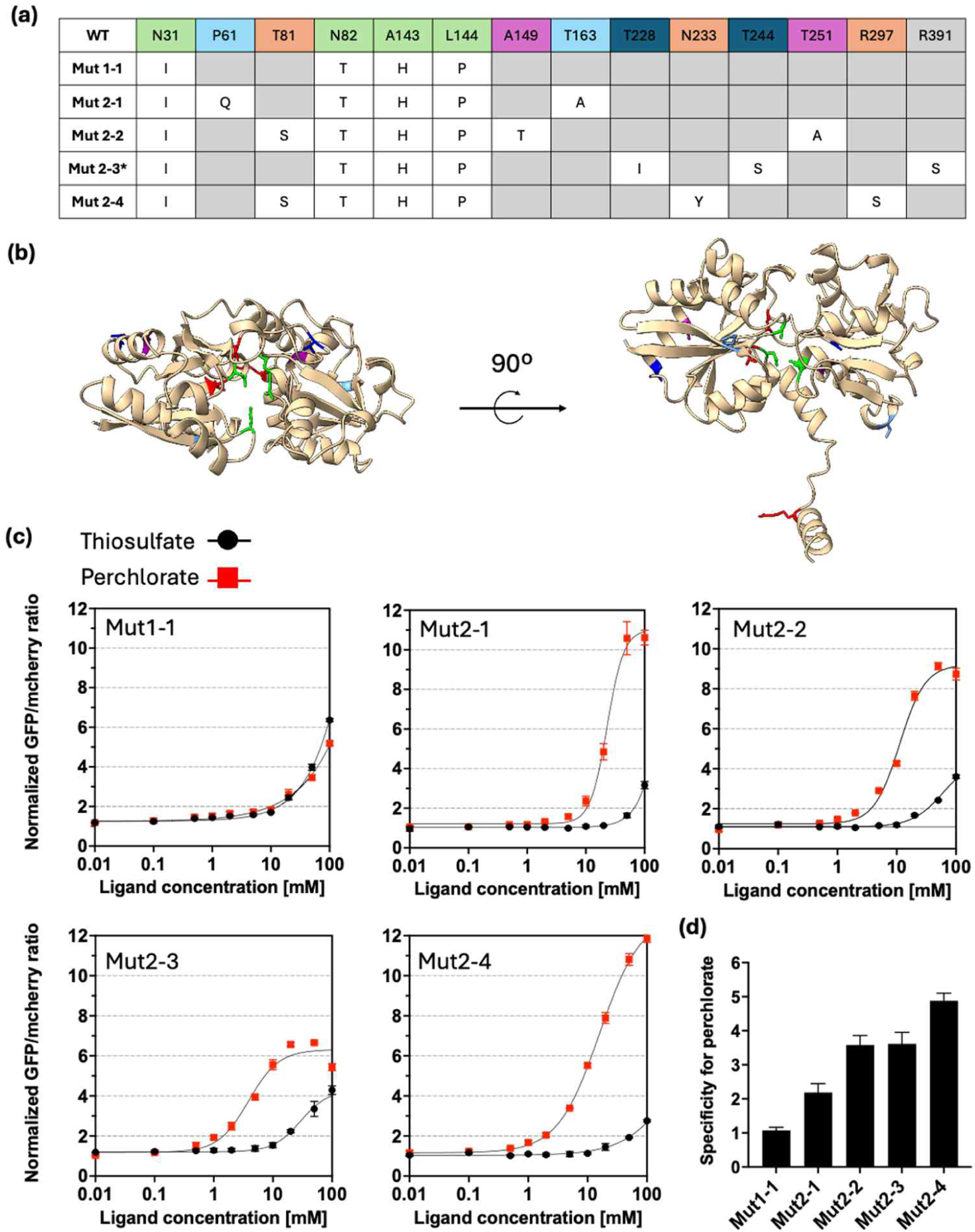
Random mutagenesis produces ThsS variants with high specificity towards perchlorate over thiosulfate. **(a)** Comparison of mutations included in each variant isolated during the second round of mutagenesis. Green color represents the original four mutations included in Mut1-1, while the other colors represent distinct mutations in each variant (Mut2-1: light blue, Mut2-2: purple, Mut2-3: dark blue, Mut2-4: orange). Note that T81S is included in both Mut2-2 and 2-4. *Mut 2-3 also contained a spontaneous mutation R391S in the cytosolic domain. **(b)** Two images depicting locations of mutated residues in Alphafold-generated model of the WT ThsS. Different colors indicate mutations included in different variant; the same color index from (a) was used. **(c)** Dose-response curves for each mutant. Compared to Mut1-1, all four variants displayed improved specificity towards perchlorate and lower detection limits. GFP/mCherry response signals were normalized with respect to the basal response of each variant without ligand addition. Each data point indicates average from three independent experiments (N=3), and the error bars indicate standard deviation. **(d)** Comparison of specificity as measured by the ratio of perchlorate-to-thiosulfate response at 10 mM ligand concentrations. Each data point indicates average from three independent experiments (N=3), and the error bars indicate standard deviation.

All four variants also showed reduced apparent sensitivity to thiosulfate relative to Mut1-1, as indicated by rightward shifts in their thiosulfate dose-response curves. Accordingly, their ratios of reporter activation by 10 mM perchlorate to that by 10 mM thiosulfate were increased (Figure 4d), demonstrating improved functional selectivity for perchlorate. Notably, the reduction in thiosulfate responsiveness emerged even though the second cycle did not include thiosulfate-dependent counterselection. Thus, stringent positive selection at low perchlorate concentrations, coupled with counterselection against basal activity, enriched variants that were both more sensitive to perchlorate and less responsive to the native ligand. Among these variants, Mut2-4 (N31I/T81S/N82T/A143H/L144P/N233Y/R297S) exhibited the lowest apparent perchlorate activation threshold, 0.31 mM, the largest inducible range, approximately 12-fold (with error range ± 0.07), and the highest perchlorate-to-thiosulfate response ratio. We therefore selected Mut2-4 as the parent for the next evolutionary cycle.

### Cycle 3: Random mutagenesis and recombination reach a performance plateau

For the third evolutionary cycle, staggered extension process (StEP) PCR (49) under error-prone conditions was used to combine recombination among all four second-round variants along with the introduction of additional point mutations. The resulting library was subjected to the same positive selection for perchlorate responsiveness and negative selection against elevated basal activity used in Cycle 2. Four variants, Mut3-1 through Mut3-4, were isolated. All four retained the original binding-pocket substitutions from Mut1-1 and contained R297S, a second-round substitution present in Mut2-4, consistent with recombination among the parental sequences. T81S and T163A were shared by Mut3-2, Mut3-3, and Mut3-4, and each variant contained no more than one additional point mutation (Figure S5a).

Despite these new combinations of mutations, none of the third-round variants show a clear functional improvement over Mut2-4. The perchlorate detection limits, dynamic ranges, and perchlorate-to-thiosulfate selectivities were, at best, comparable to those of Mut2-4 (Figure S5b). These results suggested that the improvements accessible through additional random mutagenesis and recombination had plateaued under the selection conditions used. We therefore retained Mut2-4 as the best-performing directed-evolution variant and shifted to a more targeted computational strategy for further optimization. Quantitative performance parameters for the ThsS variants isolated during each evolutionary cycle are summarized in Table 1.

**Table 1.** Specific parameters for the ThsS variants identified in this study. The parameters were obtained by the non-linear fitting of dose-response curves and subsequent calculations, as indicated in the methods section in detail. Parameters could not be calculated for Mut1-1, as the cells were killed at high perchlorate concentrations before reaching the plateau response.

| Mutation Round | Variant name | Mutations | Lower limit of detection (mM) | Upper limit of detection (mM) | Apparent K<sub>D</sub> (mM) | Dynamic Range (fold change) | Cooperativity |
|---|---|---|---|---|---|---|---|
| 1 | Mut 1-1 | N31I, N82T, A143H, L144P | NA | NA | NA | NA | 0.71 ± 0.07 |
|  | Mut 1-2 | N31R, N82R, A143I, L144R | 0.92 | 100 | 17 ± 1 | 0.77 ± 0.08 | 0.94 ± 0.23 |
| COMP 1 | G145A | N31I, N82T, A143H, L144P, G145A | 2.0 | 510 | 180 ± 90 | 4.2 ± 1.6 | 1.8 ± 0.3 |
|  | F173Q | N31I, N82T, A143H, L144P, F173Q | 1.0 | 290 | 69 ± 59 | 4.8 ± 1.8 | 1.3 ± 0.4 |
|  | L175T | N31I, N82T, A143H, L144P, L175T | 4.1 | 200 | 64 ± 25 | 3.5 ± 0.7 | 2.2 ± 0.6 |
|  | W234Q | N31I, N82T, A143H, L144P, W234Q | 4.6 | 340 | 170 ± 100 | 4.2 ± 2.5 | 1.9 ± 0.7 |
| 2 | Mut 2-1 | N31I, P61Q, N82T, A143H, L144P, T163A | 6.2 | 63 | 23 ± 1 | 9.8 ± 0.4 | 3.1 ± 0.6 |
|  | Mut 2-2 | N31I, T81S, N82T, A143H, L144P, A149T, T251A, R391S | 1.7 | 52 | 11 ± 1 | 8.0 ± 0.3 | 2.0 ± 0.2 |
|  | Mut 2-3 | N31I, N82T, A143H, L144P, T228I, T244S | 0.69 | 20 | 3.9 ± 0.5 | 5.1 ± 0.1 | 1.7 ± 0.1 |
|  | Mut 2-4 | N31I, T81S, N82T, A143H, L144P, N233Y, R297S | 0.31 | 260 | 16 ± 1 | 12 ± 1 | 1.2 ± 0.1 |
| COMP 2 | W32F | N31I, W32F, T81S, N82T, A143H, L144P, N233Y, R297S | 13 | 210 | 120 ± 50 | 4.1 ± 1.7 | 3.0 ± 1.0 |
|  | P193R | N31I, T81S, N82T, A143H, L144P, P193R, N233Y, R297S | 0.70 | 150 | 20 ± 1 | 13 ± 3 | 1.6 ± 0.2 |
|  | N233R | N31I, T81S, N82T, A143H, L144P, N233R, R297S | 0.070 | 97 | 1.8 ± 0.3 | 5.1 ± 0.5 | 1.5 ± 0.5 |
|  | P193R+N233R | N31I, T81S, N82T, A143H, L144P, P193R, N233R, R297S | 21 | 97 | 51 ± 1 | 12 ± 1 | 8.0 ± 0.3 |
| 3 | Mut 3-1 | N31I, N82T, A143H, L144P, V187A, T228I, T244S, R297S | 1.1 | 61 | 11 ± 1 | 8.7 ± 0.6 | 1.4 ± 0.1 |
|  | Mut 3-2 | N31I, T81S, N82T, R126H, A143H, L144P, T163A, R297S | 0.57 | 130 | 16 ± 3 | 11 ± 1 | 1.2 ± 0.1 |
|  | Mut 3-3 | N31I, T81S, N82T, A143H, L144P, T163A, R297S | 0.63 | 110 | 12 ± 1 | 6.3 ± 0.2 | 1.3 ± 0.1 |
|  | Mut 3-4 | N31I, T81S, N82T, A143H, L144P, T163A, L230M, R297S | 2.6 | 87 | 20 ± 2 | 8.9 ± 0.4 | 2.1 ± 0.3 |

### Computational optimization following the directed evolution plateau

Because the third cycle of directed evolution did not produce variants that outperformed Mut2-4, we returned to protCAD using experimentally validated evolved variants as starting scaffolds. Earlier molecular-dynamics simulations indicated that perchlorate did not remain in its initial modeled pose but instead moved to a different preferred location within the binding pocket. We therefore modified the protEvolver workflow to interleave sequence optimization with short MD simulations. After each design stage, the ligand and surrounding side chains were allowed to relax, and the resulting ligand position was used to guide the next round of optimization (Figure 5a).

**Figure 5.**
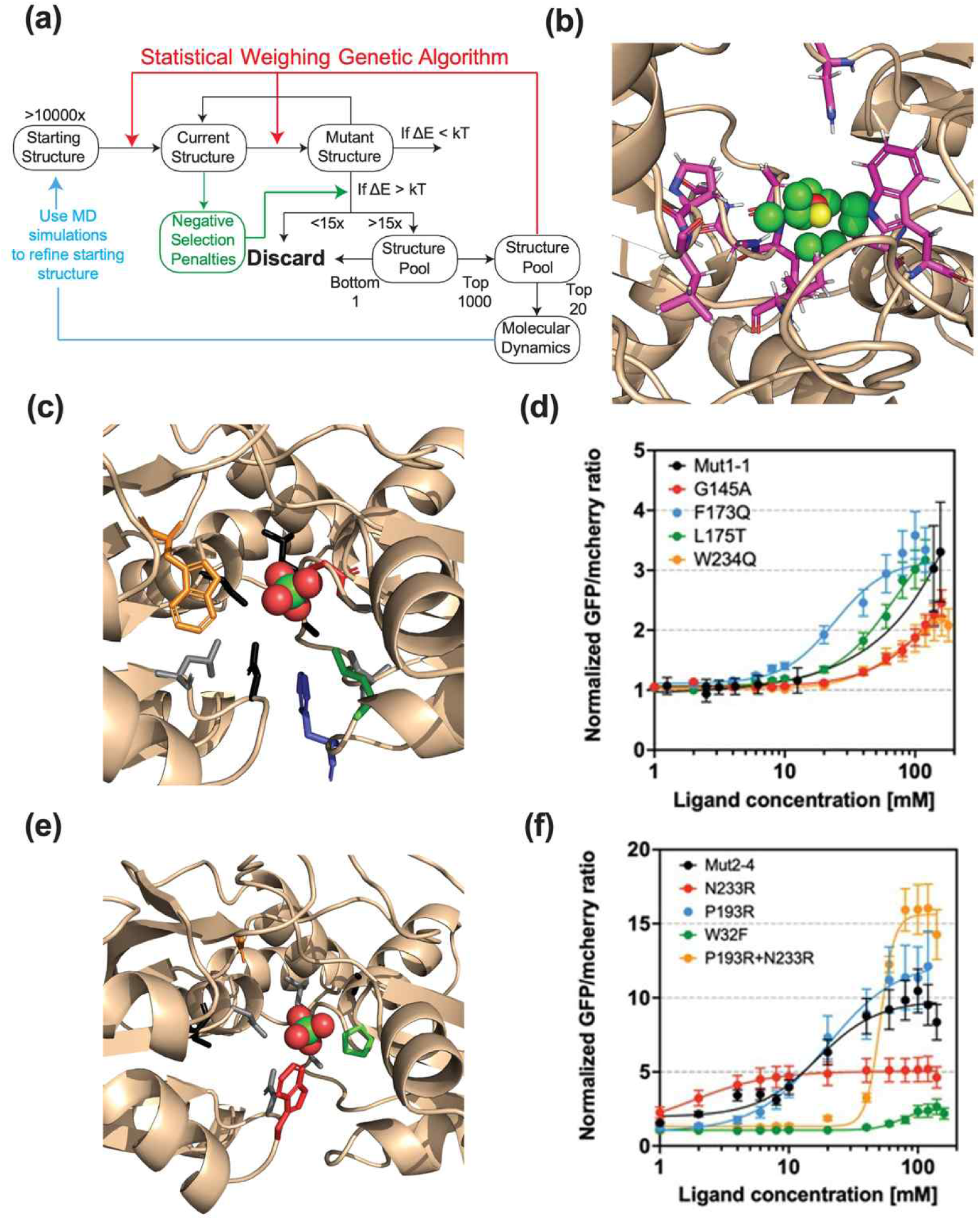
MD-guided computational optimization of evolved ThsS variants. **(a)** Modified protEvolver workflow integrating statistically weighted sequence optimization, structural minimization, energy scoring, negative-design penalties against thiosulfate recognition, and MD screening. Ligand poses sampled during MD were used to refine the starting structure for subsequent design rounds. **(b)** Superposition of perchlorate poses from MD simulations of the top-scoring Mut1-1 designs. The ThsS sensor domain is shown as a tan cartoon, variable side chains as magenta sticks, and perchlorate as spheres. The ensemble-averaged ligand position was used for the next optimization round. **(c)** Mut1-1 structural model showing the seven positions permitted to vary during the first computational campaign: residues 29, 139, 145, 146, 173, 175, and 234. Experimentally tested positions are colored to match panel D; perchlorate is shown as spheres. **(d)** Perchlorate dose-response curves for Mut1-1 and the G145A, F173Q, L175T, and W234Q variants, measured using the normalized sfGFP/mCherry fluorescence ratio. Each data point indicates average from three independent experiments (N=3), and the error bars indicate standard deviation. **(e)** Structural model of Mut2-4 showing the additional positions 32, 83, 193, and 233 permitted to vary during the second computational campaign. Positions are colored to match panel F. **(f)** Perchlorate dose-response curves for Mut2-4 and variants containing the indicated individual substitutions. Each data point indicates average from three independent experiments (N=3), and the error bars indicate standard deviation.

We first evaluated this MD-guided approach using Mut1-1. Alphafold3 (32) was used to generate a starting model of the Mut1-1 periplasmic sensor domain, comprising residues 23–309 of ThsS. Seven positions — 29, 139, 145, 146, 173, 175, and 234 — were selected for optimization because they surround the four positions randomized during Cycle 1 and were predicted to tolerate substitution without disrupting signaling (Figure 5b). Designs were generated using a new implementation of protEvolver (45) within the protCAD software package. (44, 50) Residues 29, 139, 145, 146, 173, 175 and 234, which are both immediately adjacent to the four site-saturated mutational sites and identified as residues which can be altered while retaining signal transduction were selected as targets for computational optimization (Figure 5c-d). Short MD simulations of the ten top-scoring sequences showed that perchlorate consistently shifted relative to the protein backbone (Figure S8). We therefore used the ensemble-averaged ligand position from these simulations as the starting pose for a second protEvolver optimization in which the same seven positions were allowed to vary.

The second optimization generated more than 25,000 designs, 151 of which had more favorable protCAD energy scores than the parental Mut1-1 sequence (Figure S9). At positions 29, 139, and 146, the parental amino acid was most frequently retained. At the remaining four positions, the most frequently selected substitutions were G145A, F173Q, L175T, and W234Q. Mut1-1 variants carrying each of these substitutions were generated by site-directed mutagenesis and characterized using the plate-reader assay. All four variants exhibited reporter activation at lower perchlorate concentrations than Mut1-1, with little change in the dynamic range of the sfGFP/mCherry response (Figure 5d). However, the same substitutions produced comparable increases in responsiveness to thiosulfate (Figure S10), indicating that they improved overall ligand sensitivity rather than perchlorate selectivity.

Because this initial campaign optimized the perchlorate-bound state without explicitly penalizing recognition of thiosulfate, we next added a negative-design term that disfavored sequences predicted to remain compatible with the native ligand. The revised workflow was applied to Mut2-4, the best-performing directed evolution variant. In addition to the seven positions considered in the Mut1-1 campaign, positions 32, 193, and 233 were allowed to vary (Figure 5e). The substitutions favored at the original seven positions were similar to those identified in the first campaign. Among the newly sampled positions, the most frequently selected substitutions were W32F, P193R, and replacement of residue 233 with arginine. These substitutions were introduced individually into Mut2-4 and tested experimentally. W32F abolished detectable perchlorate-dependent reporter activation, whereas P193R has little effect (Figure 5f). Substitution of residue 233 with arginine increases reporter output at 1 mM perchlorate but reduced the overall dynamic range.

These results show that the MD-guided protCAD workflow identifies mutations that improved the apparent perchlorate sensitivity of evolved ThsS variants. However, the gains observed in the tested variants were either nonselective, improving responses to both perchlorate and thiosulfate, or were accompanied by a reduction in dynamic range. Thus, none of the computationally generated variants simultaneously improved perchlorate sensitivity, ligand selectivity, and dynamic range relative to Mut2-4 (Figure 5f).

### Evaluating Biosensor Specificity Against Structurally Similar Analytes

While the workflow successfully achieved a significant reversal of ligand specificity from thiosulfate to perchlorate, characterization of sensor selectivity requires evaluation of responsiveness to a broader range of analytes. We expanded screening beyond the native and target ligands to help determine whether the change to perchlorate specificity come at the cost of increased response to off-target molecules. The response of sensors from different steps during the design process was measured in the presence of chlorate, arsenate, molybdate, bromate and sulfate (Figure 6).

**Figure 6.**
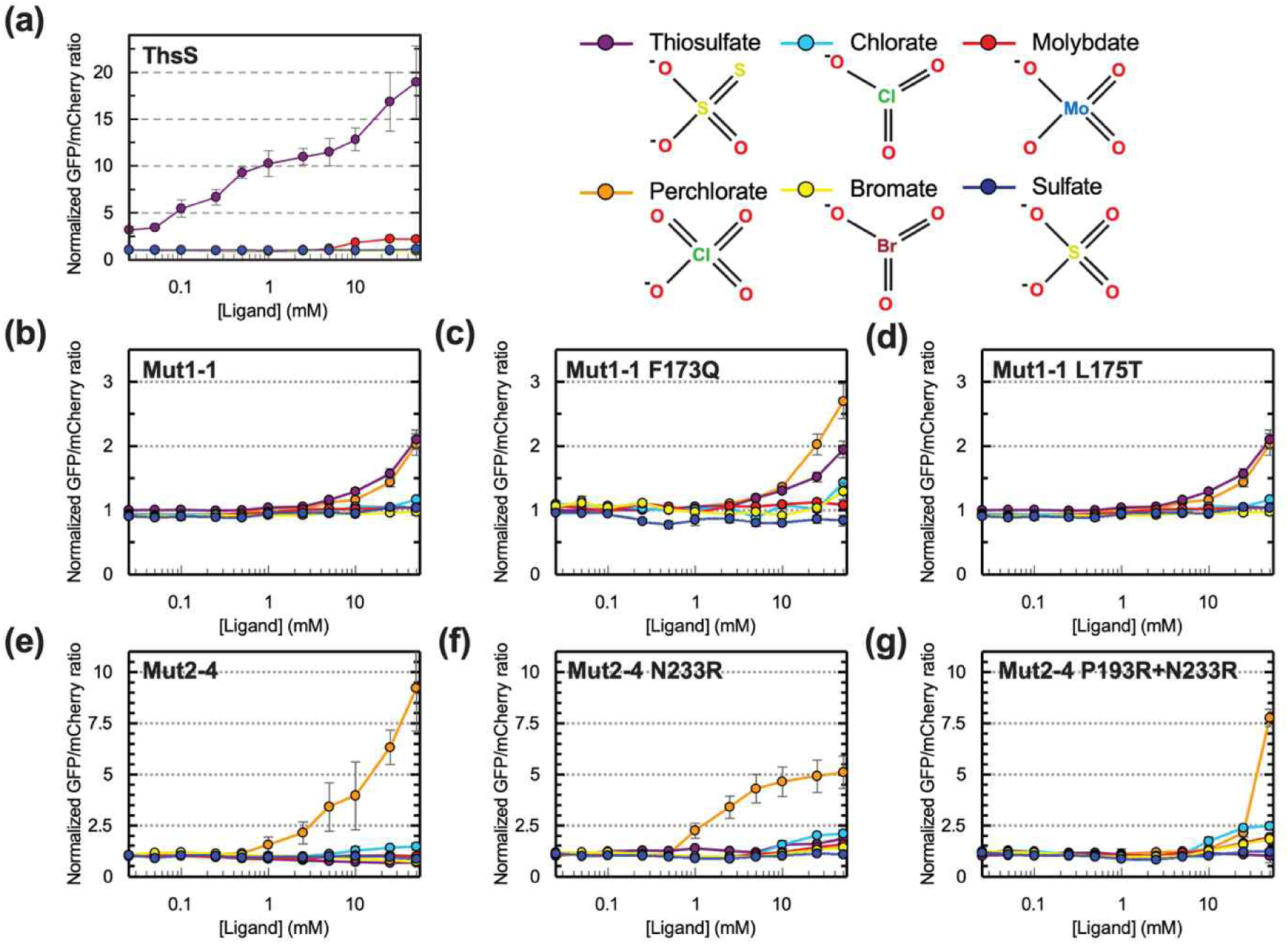
Ligand specificity of selected variants. Fluorescence intensity was measured for each mutant in the presence of perchlorate (orange), thiosulfate (purple), chlorate (cyan), molybdate (red), bromate (yellow) and sulfate (dark blue). Each data point indicates average from three independent experiments (N=3), and the error bars indicate standard deviation.

Although the WT ThsS sensor is highly specific to its native thiosulfate ligand it exhibits a weak off-target response to molybdate (Figure 6a), indicating that the naturally evolved sequence is not perfectly selective. The "generalist" variants, generated through the initial round of site-saturation mutagenesis results, showed an elevated response to both thiosulfate and perchlorate compared to all other analytes (Figure 6b-d). demonstrating that their binding pockets did not become entirely non-specific.

Subsequent rounds of error-prone PCR, which incorporated stringent negative selection, successfully yielded variants with higher selectivity for perchlorate over all other analytes (Figure 6e). Finally, while the structure-based computational design tool protCAD identified point mutations that successfully enhanced perchlorate affinity, these improvements came at the cost of some increased off-target responses (Figure 6f-g). Measurements made with arsenate or permanganate both showed ligand binding, but were complicated by toxicity and/or high absorbance in the excitation window (Figure S11).

### Biosensor Demonstration in Complex Matrices

Perchlorate can enter surface water and groundwater through releases associated with the manufacture, use, and disposal of solid rocket propellants, munitions, fireworks, and related materials. To determine whether the engineered whole-cell sensor remained functional in an environmental water matrix, we converted its output from fluorescence to bioluminescence and tested the resulting strain in laboratory medium and in nonsterile river water spiked with perchlorate. To modify the engineered perchlorate sensor for luminescence output, we replaced GFP with a *luxAB*-mVenus bioluminescence resonance energy transfer (BRET) fusion protein and also expressed *luxCDE* in the same operion (49) (Figure 7a). After these modifications, we characterized the sensitivity of the perchlorate sensor in liquid LB medium with added perchlorate. We observed a similar level of detection using the bioluminescence reporter system compared to fluorescence-based reporter, with a 1.87-fold higher luminescence (95% CI: 1.30-2.90) between the 1.56 mM perchlorate and the no perchlorate control (P = 0.013, one mean, two-tailed Student’s t-test), and a 5.24-fold higher luminescence (95% CI: 3.40-8.08) between the 3.1 mM perchlorate and the no perchlorate control (P = 0.001, one mean, two-tailed Student’s t-test) after 6 hours.

**Figure 7.**
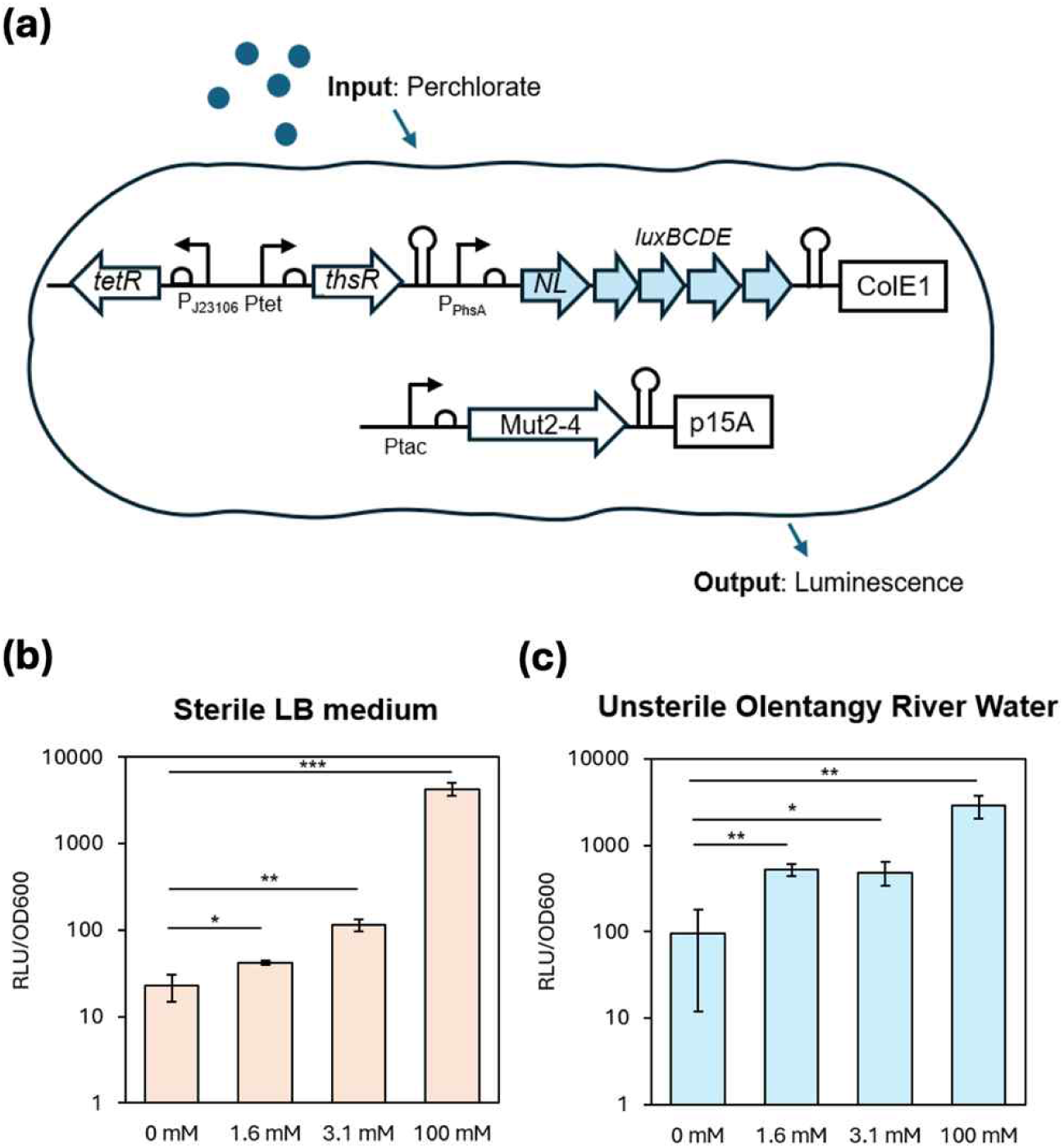
Applying the perchlorate sensing system for detecting perchlorate in environmental samples using a whole cell biosensor. **(a)** Diagram of engineered whole cell biosensor for perchlorate detection. RR = response regulator, M6 = engineered perchlorate sensing histidine kinase, NL = Nanolantern luminescence fusion protein. Origin of replication for each plasmid shown in boxes. **(b)** Performance of whole cell biosensor in sterile Luria Broth (LB) medium at different perchlorate concentrations. Briefly, the strain was grown overnight in LB medium with the appropriate antibiotics and 0.1 mM IPTG. After overnight growth, cells were washed twice in fresh LB medium, and then placed in LB medium spiked with different concentrations of sodium perchlorate at an OD600 of 0.1 and analyzed for luminescence after 6 hours. **(c)** Performance of whole cell biosensor in unsterile river water with different spiked perchlorate concentrations. Briefly, the whole cell biosensor was grown overnight in terrific broth (TB) medium, then cells were washed twice in TB medium and then inoculated in perchlorate spiked river water Strains were grown overnight in terrific broth with 0.1 mM IPTG and appropriate antibiotics. After overnight growth, strains were inoculated in unsterile, perchlorate spiked river water with 0.1 mM IPTG at an OD600 of 0.2. Samples were analyzed for luminescence after 6 hours. For (b) and (c), RLU indicates relative luminescence units, bars represent the average of three replicates (N=3), error bars are +-1 standard deviation. See Materials and Methods for more information.

Next, we tested our whole cell biosensor system in unsterile river water collected from the Olentangy River near Battelle’s Columbus, OH site. To improve response, we replaced Luria Broth (LB) with Terrific Broth (TB) to provide additional nutrients and inoculated cells into the river water with various levels of perchlorate and incubated whole cell biosensors for six hours (Figure 7b). While we observed increased background from the river water samples, we observed a 7.15-fold increase in luminescence (95% CI: 2.56-19.93) output at 1.6 mM perchlorate in unsterile river water compared to the no perchlorate control (P = 0.003, one mean, two-tailed Student’s t-test) and a 6.43-fold increase in luminescence (95% CI: 2.18-19.0) output in 3.1 mM perchlorate compared to the no perchlorate control (Figure 7c, P = 0.016, one mean, two-tailed Student’s t-test). For 100 mM perchlorate we observed a 30.6-fold increase (95% CI: 13.4-111) compared to the no perchlorate control (P = 0.004, one mean, two-tailed Student’s t-test), indicated an improved fold-change using a luminescent output compared to a fluorescent output. We attribute the improved fold-change of the biosensor in the unsterile river water conditions to incubation with TB which has additional yeast extract for protein synthesis. Overall, these results highlight the potential of this system for environmental perchlorate sensing.

## Conclusions

In this study, we reprogrammed the thiosulfate-responsive sensor histidine kinase ThsS to respond preferentially to perchlorate, an analyte for which to our knowledge no natural TCS sensor has been reported. The design pathway followed a generalist-to-specialist pattern: Site-saturation mutagenesis at four residues proximal to the predicted binding pocket generated Mut1-1 and Mut1-2, which acquired perchlorate-dependent signaling while retaining responsiveness to thiosulfate and therefore represented functionally promiscuous intermediates. Error-prone PCR across the periplasmic sensor domain, combined with positive selection for activation at low perchlorate concentrations and counterselection against elevated basal activity, subsequently produced Mut2-1 through Mut2-4. These variants showed lower apparent perchlorate activation thresholds and increased perchlorate-to-thiosulfate response ratios. This re-specialization is consistent with models of molecular evolution in which promiscuous intermediates provide access to new functional landscapes (51-53), StEP mutagenesis and recombination did not produce variants that outperformed Mut2-4, indicating that performance had plateaued under the mutational and selection conditions tested.

Computational design became particularly valuable after random evolution reached this plateau. Initial MD simulations showed that perchlorate moved away from its starting docked position, revealing a limitation of optimizing sequences against a single static structural model. We therefore interleaved protCAD sequence optimization with short MD simulations, allowing the ligand and surrounding side chains to relax between design stages and focusing subsequent optimization on the ligand position preferred by the evolving complex. When applied to Mut1-1, this MD-guided workflow identified substitutions, including F173Q and L175T, that shifted perchlorate-dependent activation toward lower concentrations. The comparable increase in thiosulfate responsiveness produced by these substitutions showed that they enhanced general ligand sensitivity rather than perchlorate selectivity. We therefore reapplied the workflow to Mut2-4 with an explicit negative-design term that disfavors retention of thiosulfate recognition—a selective pressure that was not imposed during the final directed evolution cycles. This campaign identified mutations at positions 193 and 233 that improve individual performance dimensions, including perchlorate sensitivity, signal output, and selectivity. These calculations generated experimentally validated improvements after stochastic evolution had stalled, and they expose an apparent trade-off between ligand recognition and signal transmission. Thus, computation served not merely as an initial screening tool: MD revised the structural hypothesis, positive design optimized response to the target ligand, and negative design directly addressed retention of the native ligand.

These results also illustrate why TCS reprogramming cannot be treated solely as a ligand-binding problem. SHKs can transmit signals by shifting conformational equilibria within preformed dimers, allowing mutations in the sensor domain to affect both ligand recognition and propagation of the signal through the transmembrane helices to the kinase core (54). By confining library mutagenesis to the periplasmic sensor domain, we broadly explored ligand-recognition sequence space while preserving the downstream DHp and CA domains. AlphaFold-based modeling, docking, and MD refinement favored perchlorate engagement within an internal cavity rather than at the dimer interface, supporting a working model in which evolution reshaped a pre-existing pocket rather than creating an entirely new binding mode. This architecture is consistent with the diversity of ligand-recognition mechanisms among SHK domains, in which ligands may bind near dimer interfaces, as in NarX, or within internal cavities, as in CitA and TodS (55-58). At the same time, the loss of WT sensor function caused by P193G and the distinct effects of mutations at positions 193 and 233 emphasize that reporter output depends on signal coupling as well as ligand recognition. Because the present assay measures integrated pathway activation rather than ligand binding directly, future structural and biophysical studies are needed to distinguish changes in ligand affinity from changes in signal transmission and kinase–phosphatase balance.

As an initial test of environmental applicability, we converted the sensor to a self-contained luminescent output by replacing sfGFP with a LuxAB–mVenus reporter and constitutively integrating *luxCDE*. The resulting strain retained perchlorate-dependent luminescence in nonsterile Olentangy River water spiked with perchlorate, demonstrating that the engineered signaling circuit remained functional in an environmental water matrix. We demonstrated our whole cell biosensor could detect perchlorate in unsterile environmental samples at concentrations of 1.56 mM (1,556 ug/L) which has been detected above this threshold multiple times in drinking water near ordinance and explosives facilities. While the EPA has proposed limits for perchlorate in drinking water ranging from 20 to 80 ug/L, concentrations, we believe this threshold could be addressed with additional engineering of the sensor using strategies outlined above. We foresee uses of this system for whole cell biosensing focused on environmental monitoring applications, and future work includes biocontainment strategies for whole cell biosensors along with dispersion over potential areas for environmental monitoring of perchlorate contamination which could be regularly monitored by drones equipped with optical cameras or permanent camera systems to prevent unintended release of perchlorate.

Overall, this study establishes a generalizable approach for on-demand engineering of highly specific TCS-based biosensors that integrates rational pocket mutagenesis, whole-protein growth selection, and MD-guided protCAD optimization with explicit negative design against the native ligand. The resulting perchlorate-responsive ThsSR variants demonstrate both the feasibility of assigning a new chemical input to a bacterial TCS and the broader potential of integrating computation with directed evolution to create microbial biosensors with user-defined ligand specificity. Further improvements may require engineering beyond the periplasmic binding pocket. The magnitude and duration of TCS output are determined jointly by ligand recognition, transmembrane coupling, and the autokinase and phosphatase activities of the DHp and CA domains (59). Combining sensor-domain optimization with tuning of these downstream signaling functions and broader high-throughput screening may therefore access improvements unavailable through binding-pocket redesign alone (60).

## Materials and Methods

### Plasmid construction

Plasmids PKD182 and PKD184 were obtained from Addgene (plasmids #90952 and #90953), which were kindly deposited by the Tabor research group. PKD182 was used as its original form, and PKD184 was modified to place superfolder GFP and sacB genes under P_phsA_ promoter. Each gene sequence was ordered as gBlocks fragment (Integrated DNA Technology), and cloned into the plasmid using Gibson Assembly (NEB). This modified plasmid was named PKD184-dual, and the plasmid map is included as Figure S2. Both PKD184 and PKD184-dual was then transformed into NEB5-alpha competent *E. coli* strain (NEB), further cultured and made competent again to allow for the further co-transformation with PKD182-based mutagenesis library.

### Optimization of the dual selection system

To confirm the functioning of the dual selection system, *E. coli* NEB5-alpha competent cells were co-transformed with PKD182 and pKD184 plasmids. Colony was picked and inoculated into the starter overnight culture in LB media. Subsequently, inoculated culture was diluted 100x in 100 µL LB media in a 96-well plate containing 25 µg/mL chloramphenicol, 50 µg/mL streptomycin, 0.1 mM IPTG and varying amounts of thiosulfate from 0 to 10 mM. To test the positive selection, kanamycin of varying concentration from 0 to 375 µg/mL was used, and to test the negative selection, sucrose of varying concentration from 0 to 7.5 mg/mL was used. 96-well plate was incubated with constant shaking in a microplate reader (Molecular Devices) at 37°C for 12 hours, and OD600 was measured every 10 mins to measure the cell growth.

### Site-directed mutagenesis

Alphafold3 was used to construct the structure model of ThsS protein, and Autodock Vina was used to predict the ligand binding site and nearby residues. Primers 1,2 and 3,4, which contained NNK degenerate codons (IDT), were used to amplify PCR fragments containing randomized N31, N82 and A143, L144 residues, respectively. Primers 5, 6 were used to amplify the backbone based on PKD182 plasmid. Golden Gate Assembly (NEB) was used to assemble the three fragments to yield combinations of site-saturation mutagenesis library in four different sites. All PCR amplifications were performed using Q5 Hot Start High-Fidelity PCR kit (NEB). Constructed library was purified using Zymospin PCR purification kit (Zymo Research) and transformed into competent *E. coli* cells already transformed with PKD184-dual plasmid. Resulting lawn of bacterial cells were collected and used for downstream selection process; a separate plate was transformed with diluted cells to estimate the library size by counting the number of transformants.

### Error-prone PCR mutagenesis

Primers 5 and 6 were used for the error-prone amplification of the N-terminal sensor region of ThsS gene, using GeneMorph II Random Mutagenesis Kit (Agilent). Simultaneously, primers 7 and 8 were used to amplify the backbone of PKD182 plasmid, and Golden Gate Assembly was used to assemble both fragments to build the error-prone plasmid library, which were transformed into competent E. coli cells using the same method as above site-directed mutagenesis. Staggered error-prone PCR was also performed using the same protocol, but instead of conventional PCR cycling, 80 cycles of 10 s annealing and 5 s extension was used to ensure the shuffling happens during amplification steps.

### Growth-based selection of transformed mutagenesis library

For the positive selection, *E. coli* cells containing transformed library and PKD184-dual were incubated in LB media for 12 hours in 37°C, and subsequently plated onto the LB-agar plates containing 25 µg/mL chloramphenicol, 50 µg/mL streptomycin, 375 µg/mL kanamycin, 0.1 mM IPTG and varying amounts of perchlorate from 0 to 100 mM. For the negative selection, colonies obtained from the positive selection plates were mixed and incubated in liquid LB media culture containing 25 µg/mL chloramphenicol, 50 µg/mL streptomycin, 0.1 mM IPTG and 5mg/mL sucrose. Once the culture became cloudy, cells were plated onto LB-agar plates containing 25 µg/mL chloramphenicol and 50 µg/mL streptomycin. Individual colonies were isolated and further tested for their ligand responsiveness.

### Ligand dose-response test

Starter cultures were inoculated from frozen glycerol stocks and incubated overnight in LB media containing antibiotics. The next day, inoculated seed cultures were diluted 100x into 1 mL M9 minimal media in 96 deep-well plate containing 25 µg/mL chloramphenicol, 50 µg/mL streptomycin, 0.075mM IPTG and varying concentrations of thiosulfate or perchlorate. The plate was shaken in 37°C incubator for 4 hours, then subsequently centrifuged at 4000 g for 15 min, buffer exchanged with tris buffer containing 20 mM Tris-HCl, 200 mM NaCl, pH 7.4. 200 µL of resuspended cells were transferred to black, clear bottom 96-well plate and the fluorescence was measured using microplate reader (Molecular Devices, Tecan). GFP fluorescence was measured using 460 nm excitation and 515 nm emission, and mcherry fluorescence was measured using 550 nm excitation and 610 nm emission. To construct the dose-response curve, the ratio of GFP fluorescence to mcherry fluorescence was used, and each value was further normalized with respect to the basal level GFP-to-mcherry ratio.

### Dose-response curve fitting and parameter calculation

Data were processed using Microsoft Excel and GraphPad Prism. Responds were calculated as GFP over mCherry signal ratio to normalize the expression difference between cells. The resulting ratios were further normalized through dividing by the mean blank response. Curve fitting was performed using the following embedded Prism equation:

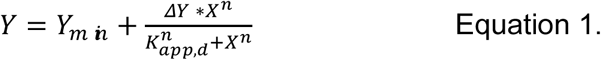

where Y represents the normalized GFP/mCherry ratio, Y_min_ is the background indicating basal expression, ΔY is calculated by Y_max_ substrates Y_min_ indicating signal amplitude, X is the ligand concentration (mM), K_app,d_ is the apparent dissociation constant (mM), and n is the Hill coefficient.

While using the Prism curving fitting on the experiment, 95% confidence interval value was also calculated for each parameter. To determine the limit of detection (LOD), where responds are statistically significantly different from blank value, the upper bound of 95% interval for Y_min_ (Y_95min_) was used in Equation 2. Similarly, the upper limit of detection (UOD) was calculated using the lower bound of the 95% confidence interval of maximum GFP/mCherry ratio (Y_95max_) (Equation 3).

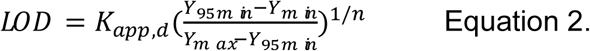

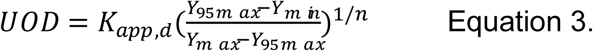

*protCAD design.* The computational design software package, protCAD (44, 50), utilizes AMBER ff14sb (61) nonbonding terms, the Dunbrack bbdep rotamer library (62) and implicit solvation using a local dielectric term (44) to score different rotamer conformations. Within protCAD, the program protEvolver (45) utilizes a genetic algorithm to bias amino acid substitutions based upon the frequency in which the appear in a lowest scoring sequence pool. The software library, including the protEvolver program, can be found at github.com/protcad/protcad. The original position of perchlorate was based upon homologous modeling with PDB ID: 7ZCK (63) and 5lV1 (64) (Figure S7). To incorporate a negative penalty for binding to off target molecules, each generation of structures attempts to lower a fitness score, given as:

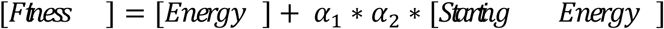

where

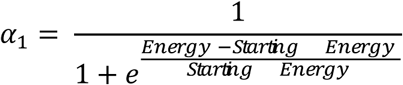

and

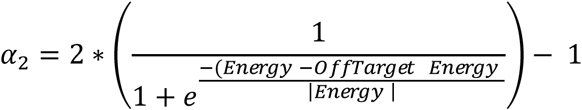

These terms are used to normalize the difference between the energy and off-target energy scores so that outliers with very large scores do not dominate the sequence pool.

### Molecular Dynamics Simulations

All simulations were performed using the OpenMM toolkit (65) with AMBER14 forcefield parameters, TIP3P-FB explicit water model, Particle Mesh Eswald periodic boundary conditions and constant pressure (T=300 K). Perchlorate parameters were obtained from the Manchester AMBER parameter database (amber.manchester.ac.uk) (66).

### Strains, Media, and Inducers

E. coli NEB 5-alpha (New England Biolabs, Ipswich, MA, USA) was used for both cloning and experimentally assaying all genetic constructs. LB Miller medium (Fisher Scientific, Waltham, MA, USA) was used for cloning. Cultures for overnights and assays were grown in LB Miller media (67) unless otherwise specified. Terrific broth was used for growth in the assay for perchlorate detection in river water. River water was taken from the Olentangy river (Columbus, Ohio). Antibiotics used for selection were 50 μg/mL kanamycin (GoldBio, St. Louis, MO, USA), 100 μg/mL spectinomycin (GoldBio), and 25 μg/mL chloramphenicol (GoldBio). Inducers used were sodium perchlorate (Sigma-Aldrich) and Isopropyl β-d-1-thiogalactopyranoside (IPTG; Sigma-Aldrich). IPTG was used at a concentration of 0.1 millimolar in each experiment.

### Perchlorate kinetic assays

Strains were grown overnight with shaking at 37°C in LB media with appropriate antibiotics. After overnight growth strains were diluted into a 96-well plate to an OD_600_ of 0.1 in fresh LB spiked with sodium perchlorate, antibiotics, and 0.1 mM IPTG. The total volume in each well was 200 µL. The plate was placed into a (Insert plate reader) and grown at 37°C with orbital shaking (2 mm amplitude). OD_600_ and total luminescence were measured at set intervals.

### Detection of perchlorate in river water

Strains were grown overnight in TB with 0.1 mM IPTG and appropriate antibiotics. After overnight growth, samples were diluted to an OD_600_ of 2.0 in fresh media with appropriate antibiotics. Strains were further diluted into a 96-well plate with river water spiked with sodium perchlorate, antibiotics, and 0.1 mM IPTG to a final OD_600_ of 0.5 or 0.2. Overnights were also mixed with spiked river water in a ratio of 20% overnight culture and 80% spiked river water. The total volume in each well was 200 µL. The plate was placed into a (Insert plate reader) and grown at 37°C with orbital shaking (2 mm amplitude). OD_600_ and total luminescence were measured using a TECAN M200 microplate reader. Confidence intervals were calculated using a 95% confidence level and values were calculated on the log scale and back-transformed to the linear scale.

## Supporting information

Supplementary Information for Lim et al

## Acknowledgements

This work was supported by DARPA’s TELLUS program (HR001124C0441). The views, opinions and/or findings expressed are those of the author and should not be interpreted as representing the official views or policies of the Department of Defense or the U.S. Government.

