## Supplementary Information for Lim et al for "Reprogramming a thiosulfate bacterial two-component system into a perchlorate biosensor"

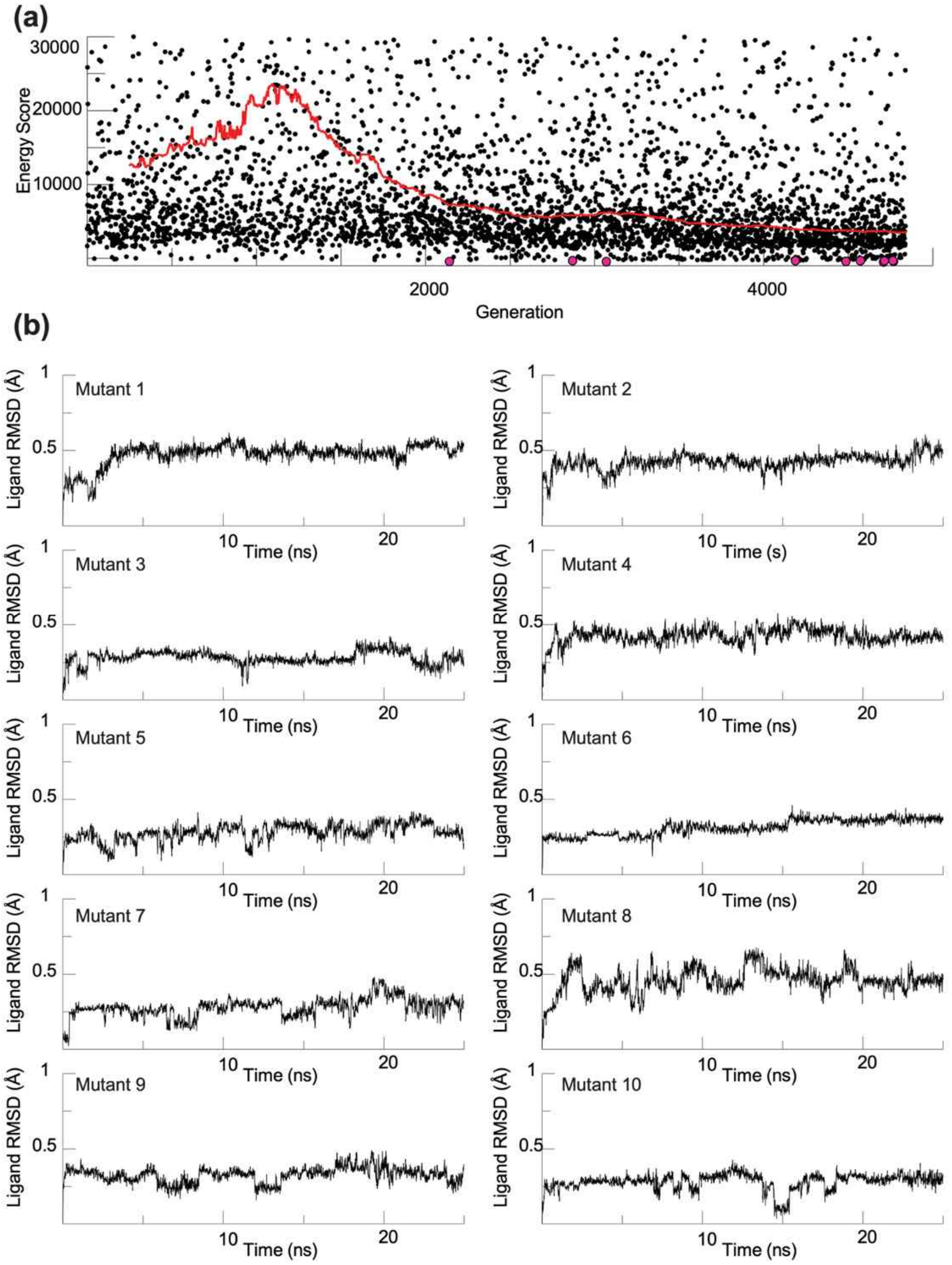

**Figure S1.** protCAD redesign of the ThsS binding pocket for perchlorate (a) protCAD energy score of every sequence generated during the initial design campaign against WT ThsS, plotted by generation. Each black point is one candidate sequence. The red trace is the running median energy score over a 500-generation window. Magenta points mark the ten lowest-scoring sequences that were carried forward. (b) Ligand RMSD of perchlorate relative to the protein backbone during MD simulations of each of the ten designs. All ten retain the ligand within the pocket, although in several designs the ligand settles into a pose displaced from the modeled starting position.

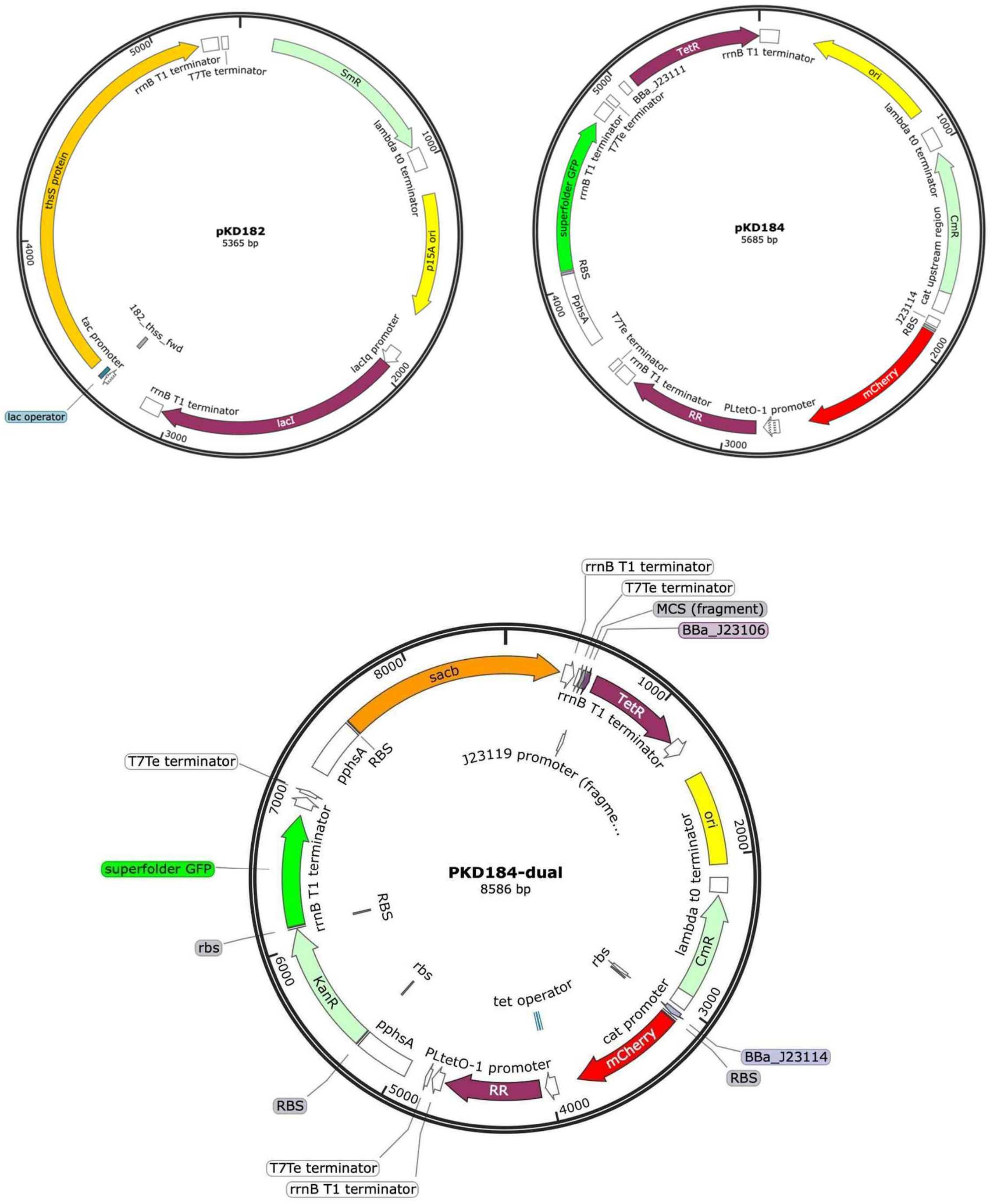

**Figure S2.** Plasmid maps of the PKD182 (Addgene #90952), PKD184 (Addgene #90953), and PKD184-dual plasmids constructed in this study. The plasmid PKD194-dual was modified based on the PKD184 plasmid.

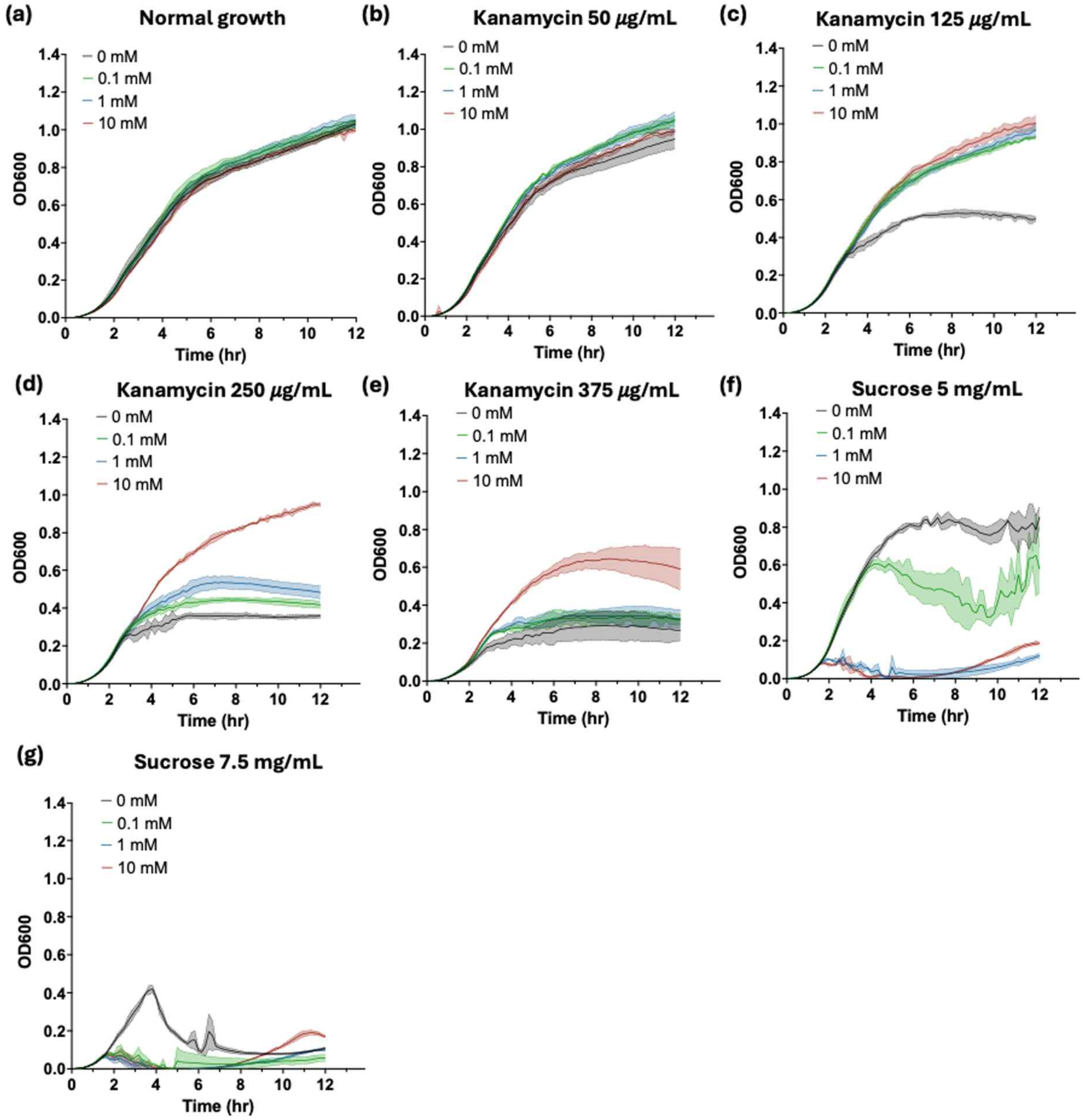

**Figure S3.** Growth curves of *E. coli* cells containing the dual-selection system, cultured under (a) normal condition, (b – e) varying kanamycin concentrations of 50, 125, 250, 375 µg/mL, (f – g) varying sucrose concentrations of 5, 7.5 mg/mL. Different color indicates different concentrations of thiosulfate (black: 0 mM, green: 0.1 mM, blue: 1 mM, red: 10 mM). Each data point indicates average from three independent experiments (N=3), and the error ranges indicate +/- one standard deviation.

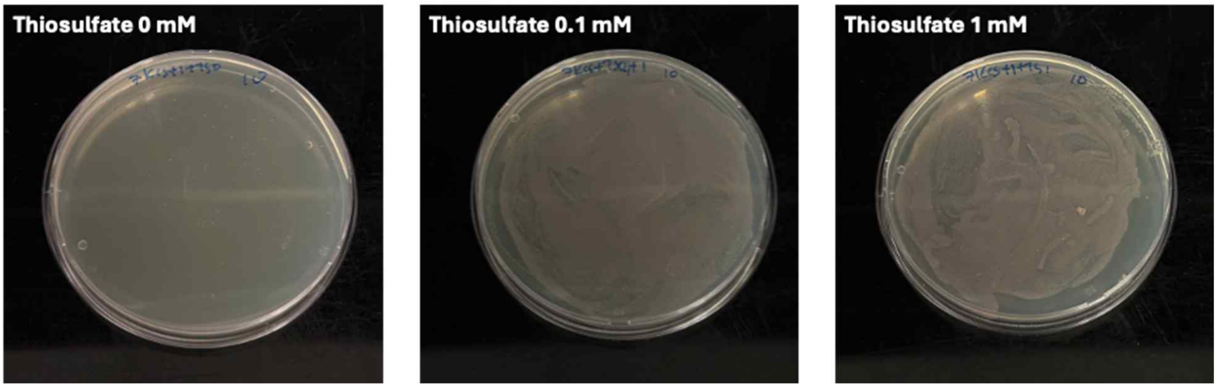

**Figure S4.** Demonstration of positive selection on solid LB-agar plates. Each plate contained 375 µg/mL kanamycin, 0.1 mM IPTG and varying amounts of thiosulfate. Clear difference in colony growth was observed from the plates containing no thiosulfate and 0.1, 1 mM of thiosulfate, indicating that solid-based plating can also be used during the positive selection cycle of directed evolution.

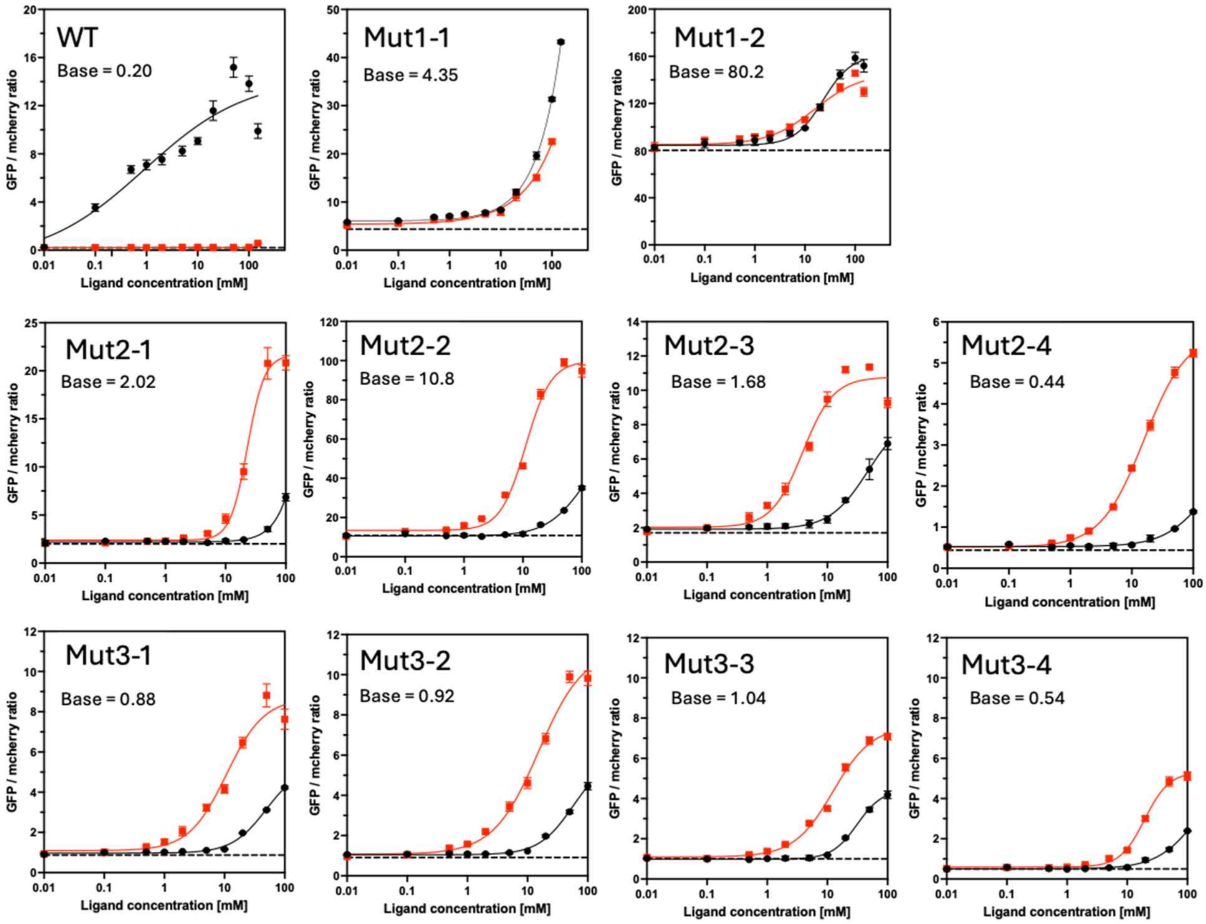

**Figure S5.** Dose-response curve for each ThsS variant without normalization. Basal response is indicated for each graph. Each data point indicates average from three independent experiments (N=3), and the error bars indicate standard deviation.

(a)

| WT | N31 | N82 | A143 | L144 | P61 | T81 | A149 | T163 | T228 | N233 | T244 | T251 | R297 |
|---|---|---|---|---|---|---|---|---|---|---|---|---|---|
| M1-1 | I | T | H | P | | | | | | | | | |
| M2-1 | I | T | H | P | Q | | | A | | | | | |
| M2-2 | I | T | H | P | | S | T | | | | | A | |
| M2-3 | I | T | H | P | | | | | I | | S | | |
| M2-4 | I | T | H | P | | S | | | | Y | | | S |

Mutation shuffle ↓

| WT | N31 | N82 | A143 | L144 | P61 | T81 | A149 | T163 | T228 | N233 | T244 | T251 | R297 | R126 | V187 | L230 |
|---|---|---|---|---|---|---|---|---|---|---|---|---|---|---|---|---|
| M3-1 | I | T | H | P | | | | | I | | S | | S | | A | |
| M3-2 | I | T | H | P | | S | | A | | | | | S | H | | |
| M3-3 | I | T | H | P | | S | | A | | | | | S | | | |
| M3-4 | I | T | H | P | | S | | A | | | | | S | | | M |

(b)

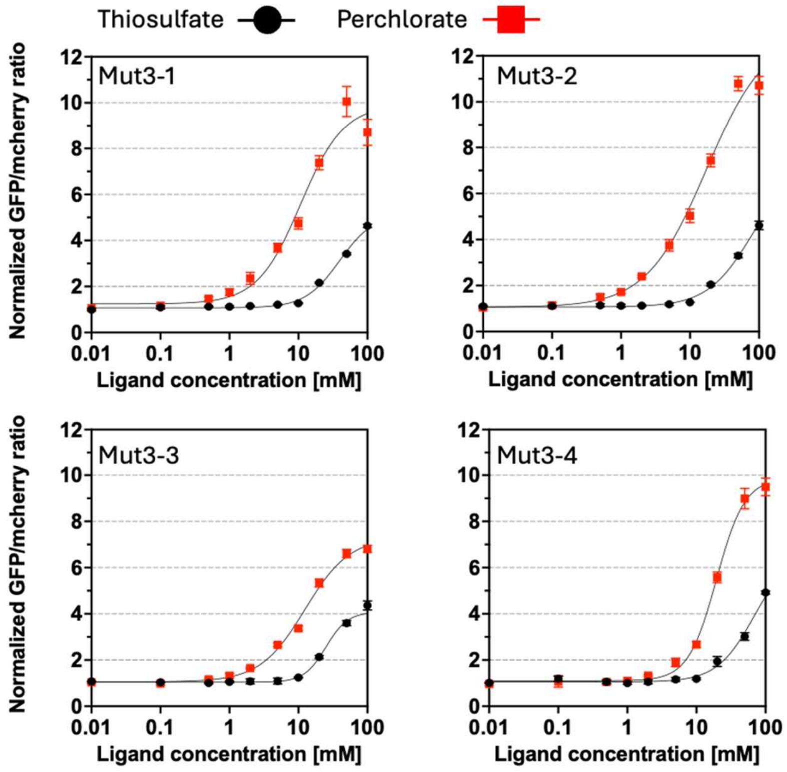

**Figure S6.** Third round of mutagenesis using staggered extension error-prone PCR. (a) Comparison of mutations included in each variant isolated during the third round of mutagenesis. Green color represents the original four mutations included in Mut1-1, orange represents mutations included in second-round variants that were shuffled, and blue represents the newly introduced mutations in the third round. (b) Dose-response curves for each mutant. GFP/mcherry response signals were normalized with respect to each variant's basal response without ligand addition. Each data point indicates average from three independent experiments (N=3), and the error bars indicate standard deviation.

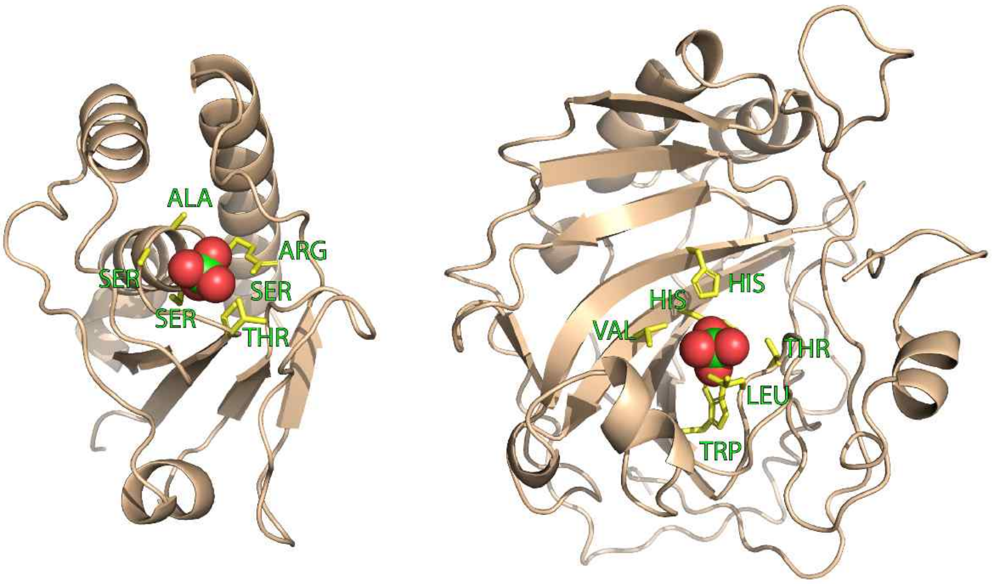

**Figure S7.** Cartoon representation of the two unique high-resolution structures in which perchlorate is specific bound. Left: PDB ID: 1JFV, Right: PDB ID: 4YGL. Perchlorate is shown in green and red.

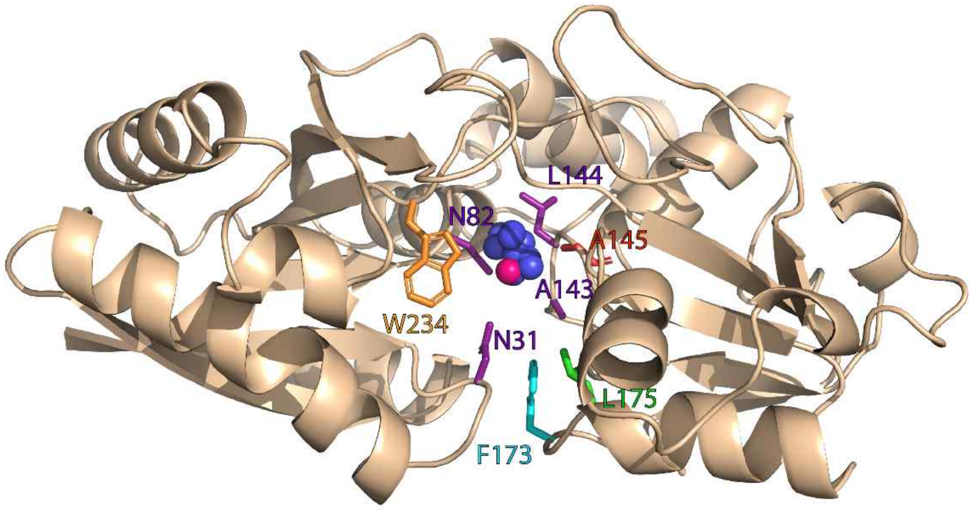

**Figure S8.** Cartoon representation of the extracellular portion of ThsS. The pink sphere represents that central chlorine atom from perchlorate, position based upon homologous structures. Blue spheres represent that average position of the perchlorate chlorine atom from molecular dynamics simulations for the top ten sequences from the initial protCAD design process.

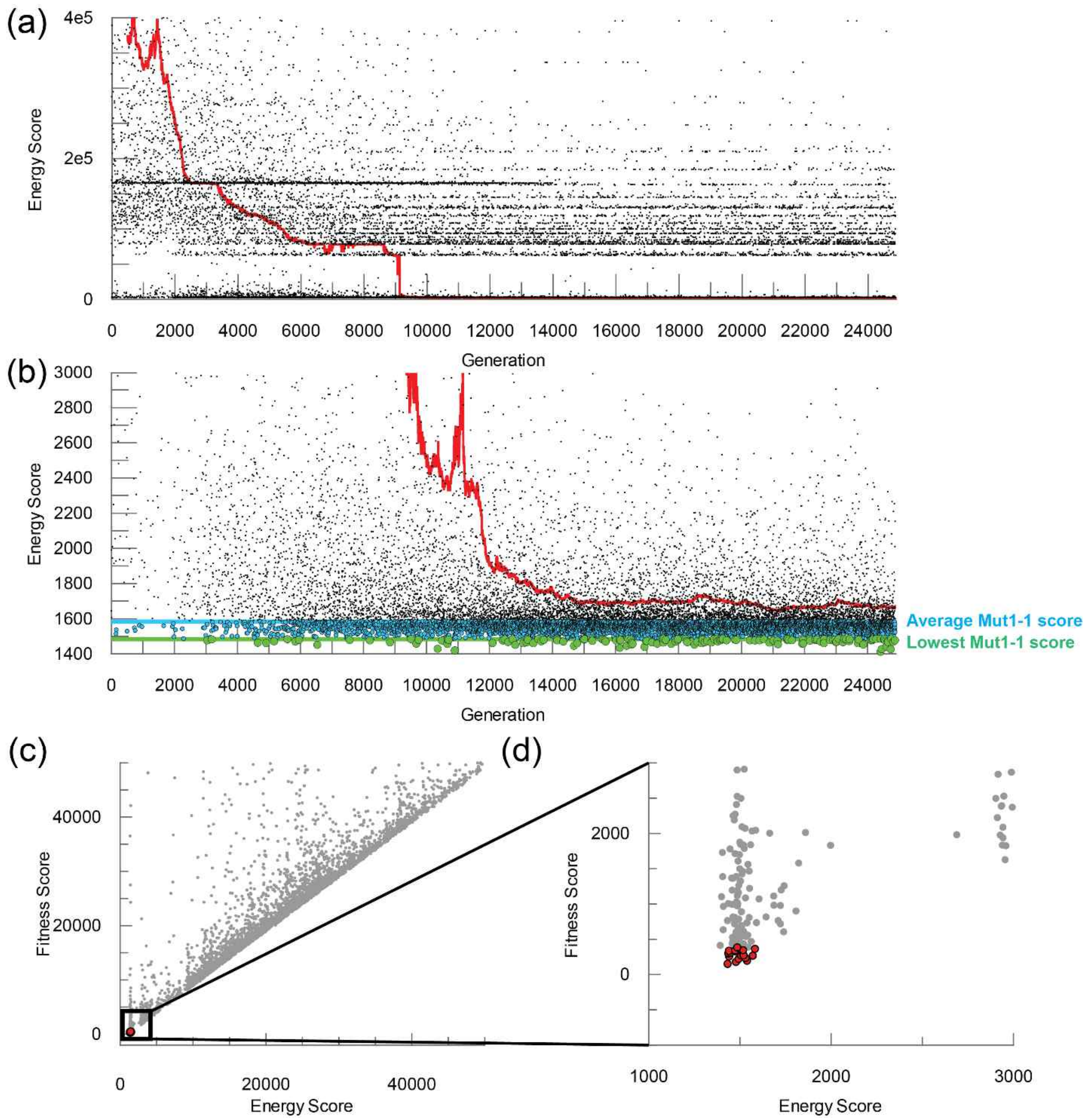

**Figure S9.** (a-b) protCAD energy scores for each generated sequence in the second round of protein design. The average and lowest energy scores from the Mut1-1 sequence in which no mutations are allowed, but rotamer rotations are permitted are shown in blue and green. The median energy score over a 500 generation window is shown in red (c-d) Fitness score versus protCAD energy score for every sequence sampled during the third round of design, which included the negative selection penalty. Top scoring sequences which were used for mutation selection are shown in magenta.

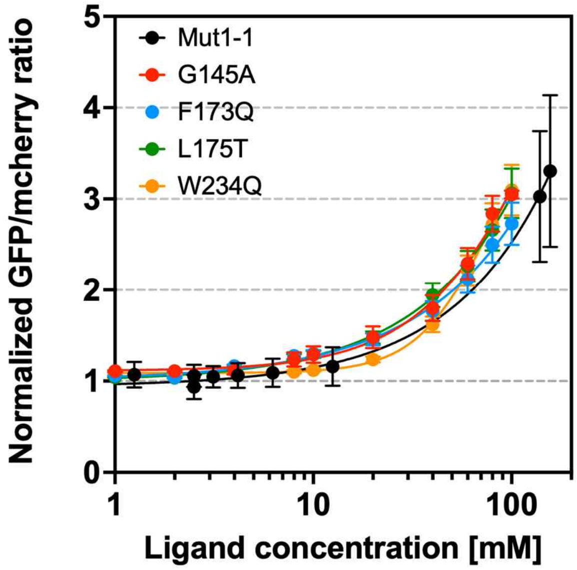

**Figure S10.** Measured thiosulfate response for the first round of protCAD mutations compared to the reference sequence, Mut1-1 (black). Each data point indicates average from three independent experiments (N=3), and the error bars indicate standard deviation.

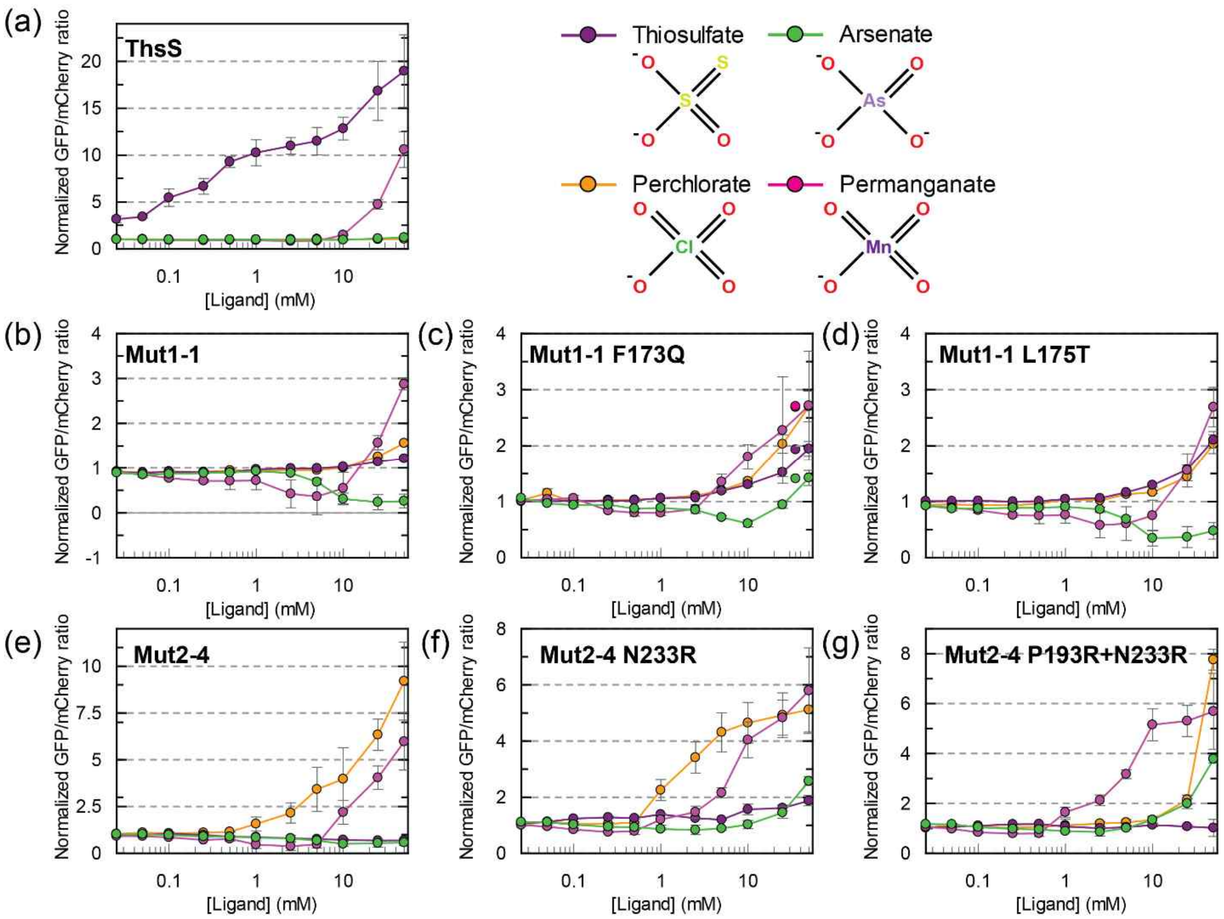

**Figure S11.** Measured response for different rounds of design when exposed to chemically-similar ligands. Fluorescence intensity was measured for each mutant in the presence of perchlorate (orange), thiosulfate (purple), arsenate (green) and permanganate (pink). Each data point indicates average from three independent experiments (N=3), and the error bars indicate standard deviation.

**Table S1.** Library generation methods and selection conditions used for the three consecutive rounds of mutagenesis performed in this study. For the positive selection, bold font represents the conditions in which colonies grew, and underlined font represents the conditions that were selected to be used for the next step.

| Rounds | Mutagenesis method | Library size | Positive selection | Negative selection |
|---|---|---|---|---|
| #1 | Site-saturation mutagenesis at 4 amino acids near predicted thiosulfate binding site | ~$10^6$ | [Kanamycin] = 375 $\mu$g/mL<br>[IPTG] = 0.1 mM<br>[Perchlorate] = 1, **<u>10</u>**, **100** mM | None |
| #2 | Random mutagenesis in ThsS extracellular domain using error-prone PCR | ~$10^6$ | [Kanamycin] = 375 $\mu$g/mL<br>[IPTG] = 0.1 mM<br>[Perchlorate] = **<u>1</u>**, **10**, **100** mM | [Sucrose] = 5 mg/mL<br>[IPTG] = 0.1 mM |
| #3 | Gene shuffling + random mutagenesis | ~$10^6$ | [Kanamycin] = 375 $\mu$g/mL<br>[IPTG] = 0.1 mM<br>[Perchlorate] = **<u>1</u>**, **10**, **100** mM | [Sucrose] = 5 mg/mL<br>[IPTG] = 0.1 mM |

**Table S2.** Primers used in this study.

| Primer number | Sequence |
|---|---|
| 1 | 5’-GGCTACGGTCTCGCGTACTGGCTNNKTGGGGTCATCAG-3’ |
| 2 | 5’-GGCTACGGTCTCATGATTGCTCAGGTACAGATATTGGCCCGGNNKAGTGATAATG-3’ |
| 3 | 5’-GGCTACGGTCTCAATCAGTACCCGCTGTCTTGGCT-3’ |
| 4 | 5’-GGCTACGGTCTCTTAGCCACCNNKNNKATGCGGGT-3’ |
| 5 | 5’-GGCTACGGTCTCTGCTACCAAGCGACCGTCGGTCTGATGCATTCCCTGGG-3’ |
| 6 | 5’-GGCTACGGTCTCGTACGCCCACATAAAACTGCTGCGGTTTAGACCACGC-3’ |
| 7 | 5’-GGCTACGGTCTCGTGGTCTAAACCGCAGCAGTTTTATG-3’ |
| 8 | 5’-GGCTACGGTCTCGTCTTTGAACAGTTTGATTACC-3’ |
| 9 | 5’-GGCTACGGTCTCGAAGAGCTGCACGTAAAAACCCCGGACTCTAGC-3’ |
| 10 | 5’-GGCTACGGTCTCGACCACGCCACAGAAGAGAAC-3’ |

**Table S3.** Sequences used in this study.

| Sequence name | DNA sequence | Reference |
|---|---|---|
| mTurquoise | atggtgagcaagggcgaggagctgttcaccggggtggtgcccatcctggtcgagc<br>tggacggcgacgtaaacggccacaagttcagcgtgtccggcgagggcgagggc<br>gatgccacctacggcaagctgaccctgaagttcatctgcacaacaggtaagctgc<br>ctgtaccatggccaactcttgtgaccactctgtcatggggtgtgcagtgcttcgcacg<br>ctacccagaccacatgaagcagcacgacttcttcaagtccgccatgcccgaaggc<br>tacgtccaggagcgcaccatcttcttcaaggacgacggcaactacaagacccgc<br>gccgaggtgaagttcgagggcgacaccctggtgaaccgcatcgagctgaaggg<br>catcgacttcaaggaggacggcaacatcctggggcacaagctggagtacaacta<br>ctttagcgacaacgtctatatcaccgccgacaagcagaagaacggcatcaaggc<br>caacttcaagatccgccacaacatcgaggacggtggtgtgcaactcgctgaccac<br>taccagcagaacactcctatcggagacggtccagtgctgctgcctgacaaccact<br>acctgagcacccagtccaagctgagcaaagaccccaacgagaagcgcgatca<br>catggtcctgctggagttcgtgaccgcagctggt | [1] |
| luxA | aaatttggaaactttttgcttacataccaacctccccaattttctcaaacagaggtaat<br>gaaacgtttggttaaattaggtcgcatctctgaggagtgtggttttgataccgtatggtt<br>actggagcatcatttcacggagtttggtttgcttggtaacccttatgtcgctgctgcatat<br>ttacttggcgcgactaaaaaattgaatgtaggaactgccgctattgttcttcccacag<br>cccatccagtacgccaacttgaagatgtgaatttattggatcaaatgtcaaaaggac<br>gatttcggtttggtatttgccgagggctttacaacaaggactttcgcgtattcggcaca<br>gatatgaataacagtcgcgccttagcggaatgctggtacgggctgataaagaatg<br>gcatgacagagggatatatggaagctgataatgaacatatcaagttccataaggta<br>aaagtaaaccccgcggcgtatagcagaggtggcgcaccggtttatgtggtggctg<br>aatcagcttcgacgactgagtgggctgctcaatttggcctaccgatgatattaagttg<br>gattataaatactaacgaaaagaaagcacaacttgagctttataatgaagtggctc<br>aagaatatgggcacgatattcataatatcgaccattgcttatcatatataacatctgta<br>gatcatgactcaattaaagcgaaagagatttgccggaaatttctggggcattggtat<br>gattcttatgtgaatgctacgactatttttgatgattcagaccaaacaagaggttatgat<br>ttcaataaagggcagtggcgtgactttgtattaaaaggacataaagatactaatcgc<br>cgtattgattacagttacgaaatcaatcccgtgggaacgccgcaggaatgtattgac<br>ataattcaaaaagacattgatgctacaggaatatcaaatatttgttgtggatttgaagc<br>taatggaacagtagacgaaattattgcttccatgaagctcttccagtctgatgtcatgc<br>catttcttaaagaaaaacaacgttcgctattatattag | [1] |
| LuxB | atgaaatttggattgttcttccttaacttcatcaattcaacaactgttcaagaacaaagt<br>atagttcgcatgcaggaaataacggagtatgttgataagttgaattttgaacagatttt<br>agtgtatgaaaatcatttttcagataatggtgttgtcggcgctcctctgactgtttctggttt<br>tctgctcggtttaacagagaaaattaaaattggttcattaaatcacatcattacaactc<br>atcatcctgtccgcatagcggaggaagcttgcttattggatcagttaagtgaaggga<br>gatttattttagggtttagtgattgcgaaaaaaaagatgaaatgcatttttttaatcgccc<br>ggttgaatatcaacagcaactatttgaagagtgttatgaaatcattaacgatgctttaa<br>caacaggctattgtaatccagataacgattttatagcttccctaaaatatctgtaaatc<br>cccatgcttatacgccaggcggacctcggaaatatgtaacagcaaccagtcatcat<br>attgttgagtgggcggccaaaaaaggtattcctctcatctttaagtgggatgattctaa<br>tgatgttagatatgaatatgctgaaagatataaagccgttgcggataaatatgacgtt<br>gacctatcagagatagaccatcagttaatgatattagttaactataacgaagatagt<br>aataaagctaaacaagagacgcgtgcatttattagtgattatgttcttgaaatgcacc<br>ctaatgaaaatttcgaaaataaacttgaagaaataattgcagaaaacgctgtcgga<br>aattatacggagtgtataactgcggctaagttggcaattgaaaagtgtggtgcgaaa<br>agtgtattgctgtcctttgaaccaatgaatgatttgatgagccaaaaaaatgtaatca<br>atattgttgatgataatattaagaagtaccacatgaaatatacctaa | [1] |
| LuxC | atgactaaaaaaatttcattcattattaacggccaggttgaaatctttcccgaaggtga<br>tgatttagtgcaatccattaattttggtgataatagtgtttacctgccaatattgaatgact<br>ctcatgtaaaaaacattattgattgtaatggaaataacgaattacggttgcataacatt<br>gtcaattttctctatacggtagggcaaagatggaaaaatgaagaatactcaagacg | [1] |

| | | |
|---|---|---|
| | caggacatacattcgtgacttaaaaaaatatatgggatattcagaagaaatggcta<br>agctagaggccaattggatatctatgattttatgttctaaaggcggcctttatgatgttgt<br>agaaaatgaacttggttctcgccatatcatggatgaatggctacctcaggatgaaag<br>ttatgttcgggcttttccgaaaggtaaatctgtacatctgttggcaggtaatgttccattat<br>ctgggatcatgtctatattacgcgcaattttaactaagaatcagtgtattataaaaaca<br>tcgtcaaccgatccttttaccgctaatgcattagcgttaagttttattgatgtagaccctta<br>atcatccgataacgcgctctttatctgttatatattggccccaccaaggtgatacatca<br>ctcgcaaaagaaattatgcaacatgcggatgttattgtcgcttggggagggccagat<br>gcgattaattgggcggtagagcatgcgccatcttatgctgatgtgattaaatttggttct<br>aaaaagagtctttgcattatcgataatcctgttgatttgacgtccgcagcgacaggtg<br>cggctcatgatgtttgtttttacgatcagcgagcttgtttttctgcccaaaacatatattac<br>atgggaaatcattatgaggaatttaagttagcgttgatagaaaaacttaatctatatgc<br>gcatatattaccgaatgccaaaaaagattttgatgaaaaggcggcctattctttagttc<br>aaaaagaaagcttgtttgctggattaaaagtagaggtggatattcatcaacgttggat<br>gattattgagtcaaatgcaggtgtggaatttaatcaaccacttggcagatgtgtgtac<br>cttcatcacgtcgataatattgagcaaatattgccttatgttcaaaaaaataagacgc<br>aaaccatatctatttttccttgggagtcatcatttaaatatcgagatgcgttagcattaaa<br>aggtgcggaaaggattgtagaagcaggaatgaataacatatttcgagttggtggat<br>ctcatgacggaatgagaccgttgcaacgattagtgacatatatttctcatgaaaggc<br>catctaactatacggctaaggatgttgcggttgaaatagaacagactcgattcctgg<br>aagaagataagttccttgtatttgtcccataa | |
| LuxD | atggaaaatgaatcaaaatataaaaccatcgaccacgttatttgtgttgaaggaaat<br>aagaaaattcatgtttgggaaacgctgccagaagaaaacagcccaaagagaaa<br>gaatgccattattattgcgtctggttttgcccgcaggatggatcattttgctggtctggcg<br>gaatatttatcgcggaatggatttcatgtgatccgctatgattcgcttcaccacgttgga<br>ttgagttcagggacaattgatgaatttacaatgtctataggaaagcagagcttgttag<br>cagtggttgattggttaactacacgaaaaataaataacttcggtatgttggcttcaag<br>cttatctgcgcggatagcttatgcaagcctatctgaaatcaatgcttcgtttttaatcacc<br>gcagtcggtgttgttaacttaagatattctcttgaaagagctttagggtttgattatctca<br>gtctacccattaatgaattgccgaataatctagattttgaaggccataaattgggtgct<br>gaagtctttgcgagagattgtcttgattttggttgggaagatttagcttctacaattaata<br>acatgatgtatcttgatataccgtttattgcttttactgcaaataacgataattgggtcaa<br>gcaagatgaagttatcacattgttatcaaatattcgtagtaatcgatgcaagatatatt<br>ctttgttaggaagttcgcatgacttgagtgaaaatttagtggccctgcgcaatttttatca<br>atcggttacgaaagccgctatcgcgatggataatgatcatctggatattgatgttgat<br>attactgaaccgtcatttgaacatttaactattgcgacagtcaatgaacgccgaatga<br>gaattgagattgaaaatcaagcaatttctctgtcttaa | [1] |
| LuxE | atgacttcatatgttgataaacaagaaattacagcaagctcagaaattgatgatttga<br>tttttcgagcgatccattagtgtggtcttacgacgagcaggaaaaaatcagaaaga<br>aacttgtgcttgatgcatttcgtaatcattataaacattgtcgagaatatcgtcactactg<br>tcaggcacacaaagtagatgacaatattacggaaattgatgacatacctgtattccc<br>aacatcggtttttaagtttactcgcttattaacttctcaggaaaacgagattgaaagttg<br>gtttaccagtagcggcacgaatggtttaaaaagtcaggtggcgcgtgacagattaa<br>gtattgagagactcttaggctctgtgagttatggcatgaaatatgttggtagttggtttga<br>tcatcaaatagaattagtcaatttgggaccagatagatttaatgctcataatatttggttt<br>aaatatgttatgagtttggtggaattgttatatcctacgacatttaccgtaacagaaga<br>acgaatagattttgttaaaacattgaatagtcttgaacgaataaaaaatcaagggaa<br>agatctttgtcttattggttcgccatactttatttatttactctgccattatatgaaagataaa<br>aaaatctcattttctggagataaaagcctttatatcataaccggaggcggctggaaa<br>agttacgaaaaagaatctctgaaacgtgatgatttcaatcatcttttatttgatactttca<br>atctcagtgatattagtcagatccgagatatatttaatcaagctgaactcaacacttgtt<br>tctttgaggatgaaatgcagcgtaaacatgttccgccgtgggtatatgcgcgagcgc<br>ttgatcctgaaacgttgaaacctgtacctgatggaacgccggggttgatgagttatat<br>ggatgcgtcagcaaccagttatccagcatttattgttaccgatgatgtcgggataatta<br>gcagagaatatggtaagtatcccggcgtgctcgttgaaattttacgtcgcgtcaatac<br>gaggacgcagaaagggtgtgctttaagcttaaccgaagcgtttgatagttga | [1] |

| | | |
|---|---|---|
| TetR | ttaagacccactttcacatttaagttgtttttctaatccgcatatgatcaattcaaggccg<br>aataagaaggctggctctgcaccttggtgatcaaataattcgatagcttgtcgtaata<br>atggcggcatactatcagtagtaggtgtttccctttcttctttagcgacttgatgctcttga<br>tcttccaatacgcaacctaaagtaaaatgccccactgcgctgagtgcatataatgca<br>ttctctagtgaaaaaccttgttggcataaaaaggctaattgattttcgagagtttcatac<br>tgtttttctgtaggccgtgtacctaaatgtacttttgctccatcgcgatgacttagtaaag<br>cacatctaaaacttttagcgttattacgtaaaaaatcttgccagctttccccttctaaag<br>ggcaaaagtgagtatggtgcctatctaacatctcaatggctaaggcgtcgagcaaa<br>gcccgcttattttttacatgccaatacaatgtaggctgctctacacctagcttctgggcg<br>agtttacgggttgttaaaccttcgattccgacctcattaagcagctctaatgcgctgtta<br>atcactttacttttatctaaacgagacatcat | [2] |
| LacI | gtgaaaccagtaacgttatacgatgtcgcagagtatgccggtgtctcttatcagacc<br>gtttcccgcgtggtgaaccaggccagccacgtttctgcgaaaacgcgggaaaaag<br>tggaagcggcgatggcggagctgaattacattcccaaccgcgtggcacaacaact<br>ggcgggcaaacagtcgttgctgattggcgttgccacctccagtctggccctgcacg<br>cgccgtcgcaaattgtcgcggcgattaaatctcgcgccgatcaactgggtgccagc<br>gtggtggtgtcgatggtagaacgaagcggcgtcgaagcctgtaaagcggcggtgc<br>acaatcttctcgcgcaacgcgtcagtgggctgatcattaactatccgctggatgacc<br>aggatgccattgctgtggaagctgcctgcactaatgttccggcgttatttcttgatgtct<br>ctgaccagacacccatcaacagtattattttctcccatgaagacggtacgcgactgg<br>gcgtggagcatctggtcgcattgggtcaccagcaaatcgcgctgttagcgggccca<br>ttaagttctgtctcggcgcgtctgcgtctggctggctggcataaatatctcactcgcaat<br>caaattcagccgatagcggaacgggaaggcgactggagtgccatgtccggttttc<br>aacaaaccatgcaaatgctgaatgagggcatcgttcccactgcgatgctggttgcc<br>aacgatcagatggcgctgggcgcaatgcgcgccattaccgagtccgggctgcgc<br>gttggtgcggatatctcggtagtgggatacgacgataccgaagacagctcatgttat<br>atcccgccgttaaccaccatcaaacaggattttcgcctgctggggcaaaccagcgt<br>ggaccgcttgctgcaactctctcagggccaggcggtgaagggcaatcagctgttgc<br>ccgtctcactggtgaaaagaaaaaccaccctggcgcccaatacgcaaaccgcct<br>ctccccgcgcgttggccgattcattaatgcagctggcacgacaggtttcccgactgg<br>aaagcgggcagtga | [2] |
| ThsR | atgcagcagcaaatcaacggcccggtctacctggtggatgatgatgaagccattat<br>cgactccatcgatttttgatggagggctacggttacaaactgaactcgtttaactgcg<br>gcgatcgctttttggcagaagtcgatctcacccaggcaggatgtgtaattctggatgc<br>gcgtatgccaggcttaactggtcctcaggtgcaacagctgctgagcgacgcgaaa<br>agcccgcttgcggtcatcttcctgaccggccatggcgatgttccgatggcggttgatg<br>cgttcaaaaatggcgcgttcgatttctttcaaaaacctgtgccgggtagcttgctcagt<br>cagtcaattgccaaaggcttgacttattcaatcgatcaacatctgaaacgtactaacc<br>aagcgttaatcgacacgctctcggaacgcgaagctcaaatttttcaactggtgattg<br>caggcaacaccaacaaacagatggctaacgagctttgcgtggctattcgtaccatt<br>gaggttcaccgtagcaaactgatgaccaaactgggtgttaacaacctggctgaact<br>ggttaaactggcgccgctgctggcacataaatccgaataa | [2] |
| ThsS | atgtcccgcctgctgctgtgtatctgtgttctgctgttctcttctgtggcgtggtctaaacc<br>gcagcagttttatgtgggcgtactggctaactggggtcatcagcaagccgttgaacg<br>ttggaccccgatgatggagtatctgaacgaacatgtgccggacgcggaatttcacg<br>tctacccgggcaacttcaaagcactgaacctggcaatggaactgggccagattca<br>gttcattatcactaacccgggccaatatctgtacctgagcaatcagtacccgctgtctt<br>ggctggcgaccatgcgttctaagcgtcacgatggtaccacttctgcgatcggttccg<br>ccattattgtccgcgcggacagcgactaccgcaccctgtacgacctgaaaggtaa<br>agtggtggctgcgtccgacccgcatgctctgggtggctaccaagcgaccgtcggtc<br>tgatgcattccctgggcatggatccggacaccttcttcggtgaaaccaagtttctggg<br>ctttccactggatccgctgctgtaccaagttcgtgatggcaacgttgacgcggccatt<br>accccactgtgcactctggaggacatggttgcacgcggcgtactgaaatcttccgat<br>tttcgtgtgctgaaccctagccgcccggatggtgtagaatgccagtgctctaccacc<br>ctgtacccgaactggtctttcgctgcgactgagtctgtatccaccgaactgtctaaag<br>aaatcacgcaggcactgctggaactgccatccgacagcccggcagctatcaaag | [2] |

| | | |
|---|---|---|
| | cgcaactgaccggctggaccagcccgatctcccaactggcggtaatcaaactgttc aaagagctgcacgtaaaaaccccggactctagccgttgggaagccgttaagaag tggctggaagaaaaccgtcactggggtatcctgtctgttctggtgttcatcattgcaac gctgtatcacctgtggattgaataccgcttccaccaaaaaagctcttctctgatcgaat ctgaacgtcagctgaaacagcaagctgttgccctggaacgtctgcaatctgctagc atcgttggtgaaattggtgcgggtctggcccacgagattaatcagccgatcgctgca attacctcttattctgaaggtggcatcatgcgcctgcaaggtaaagaacaggcggat acggatagctgcatcgaactgctggaaaaaatccacaaacagagcactcgcgca ggcgaagtggtgcaccgcatccgtggtctgctgaaacgtcgtgaagcggtgatggt agatgttaacatcctgaccctggtggaagaatccatcagcctgctgcgtctggagct ggcacgtcgcgaaatccagatcaacactcagatcaaaggtgaaccgttcttcatta ctgccgaccgcgttggcctgctgcaagttctgattaacctgatcaaaaactccctgg acgcgatcgctgaatctgataatgcccgttctggtaaaatcaacatcgaactggact ttaaagagtaccaggtaaacgtctccatcatcgataacggtccgggcctggcgatg gattctgacactctgatggctacgttttacactaccaaaatggatggcctgggcctgg gtctggcaatctgccgcgaagttatcagcaaccacgacggccacttcctgctgtcca accgtgacgacggcgttctgggctgtgtggcaaccctgaatctgaaaaaacgcgg ttctgaagtgccgatcgaagtctaa | |
| PTac | ttgacaattaatcatcggctcgtataatgtgtggaattgtgagcggataacaa | [2] |
| PLtetO-1 | tccctatcagtgatagagattgacatccctatcagtgatagagatactgagcacatc agcaggacgcactgacc | [2] |
| PphsA | ttcaagcattattatgctgtttttgaagtgaatgtgcggccatctagccgcacattttgc atctaaaacatgcagtcatcagcaaaataataaacttttccccaatatgtggtttacc acaatttacaggaattcactcctgtggtggtgcaaatttgaactgtgaattgcttcaca aacgccgctatcgcaatgtcagtatgtggtttaccacaatatctaatatcactctgctc aataacaatgatgaaaaccttaggaagaagttaattgtgttaaacagttaactaggg gctttatctaacgctctcctaaggacaactgtcattgggagatttaac | [2] |
| pKD184 | tagtgatcttatttcattatggtgaaagttggaacctcttacgggccggccaacgtctc attttcgccagatatcgatttatggctagctcagtcctaggtacaatgctagcgaatcc attaaagaggagaaaggtacccatggtgagcaagggcgaggaggataacatgg ccatcatcaaggagttcatgcgcttcaaggtgcacatggagggctccgtgaacggc cacgagttcgagatcgagggcgagggcgagggccgcccctacgagggcaccc agaccgccaagctgaaggtgaccaagggtggcccccctgcccttcgcctgggaca tcctgtcccctcagttcatgtacggctccaaggcctacgtgaagcaccccgccgac atccccgactacttgaagctgtccttccccgagggcttcaagtgggagcgcgtgatg aacttcgaggacggcggcgtggtgaccgtgacccaggactcctccctgcaggac ggcgagttcatctacaaggtgaagctgcgcggcaccaacttcccctccgacggcc ccgtaatgcagaagaagaccatgggctgggaggcctcctccgagcggatgtacc ccgaggacggcgccctgaagggcgagatcaagcagaggctgaagctgaagga cggcggccactacgacgctgaggtcaagaccacctacaaggccaagaagccc gtgcagctgcccggcgcctacaacgtcaacatcaagttggacatcacctcccaca acgaggactacaccatcgtggaacagtacgaacgcgccgagggccgccactcc accggcggcatggacgagctgtacaagtaagctttctaactaaaaaggcctccca aatcgggggggcctttttttattgataacaaaacgctcgtctaagaaaccattattatcat gacattaacctataaaaataggcgtatcacgaggccctttcgtcttcacctcgagtcc ctatcagtgatagagattgacatccctatcagtgatagagatactgagcacatcagc aggacgcactgacctatttagtttacttattaatttattaaaattatatgcagcagcaa atcaacggcccggtctacctggtggatgatgatgaagccattatcgactccatcgatt ttttgatggagggctacggttacaaactgaactcgtttaactgcggcgatcgctttttgg cagaagtcgatctcacccaggcaggatgtgtaattctggatgcgcgtatgccaggc ttaactggtcctcaggtgcaacagctgctgagcgacgcgaaaagcccgcttgcggt catcttcctgaccggccatggcgatgttccgatggcggttgatgcgttcaaaaatggc gcgttcgatttctttcaaaaacctgtgccgggtagcttgctcagtcagtcaattgccaa aggcttgacttattcaatcgatcaacatctgaaacgtactaaccaagcgttaatcga cacgctctcggaacgcgaagctcaaatttttcaactggtgattgcaggcaacacca acaaacagatggctaacgagctttgcgtggctattcgtaccattgaggttcaccgta | [2] |

| | |
|---|---|
| | gcaaactgatgaccaaactgggtgttaacaacctggctgaactggttaaactggcg<br>ccgctgctggcacataaatccgaataataatctagaccaggcatcaaataaaacg<br>aaaggctcagtcgaaagactgggcctttcgttttatctgttgtttgtcggtgaacgctct<br>ctactagagtcacactggctcaccttcgggtgggcctttctgcgtttatagtaaatcact<br>gcataattcgtgtcgctcaaggcgcactcccgttctggataatgttttttgcgccgacat<br>cataacggttctggcaaatattctgaaatgagctggttagctagtcattcaagcattatt<br>atgctgtttttgaagtgaatgtgcggccatctagccgcacattttgcatctaaaacatg<br>cagtcatcagcaaaataataaacttttccccaatatgtggtttaccacaatttacagg<br>aattcactcctgtggtggtgcaaatttgaactgtgaattgcttcacaaacgccgctatc<br>gcaatgtcagtatgtggtttaccacaatatctaatatcactctgctcaataacaatgat<br>gaaaaccttaggaagaagttaattgtgttaaacagttaactaggggctttatctaacg<br>ctctcctaaggacaactgtcattgggagatttaacaaagaggagaaatactagatg<br>cgtaaaggcgaagagctgttcactggtgtcgtccctattctggtggaactggatggtg<br>atgtcaacggtcataagttttccgtgcgtggcgagggtgaaggtgacgcaactaatg<br>gtaaactgacgctgaagttcatctgtactactggtaaactgccggtaccttggccgac<br>tctggtaacgacgctgacttatggtgttcagtgctttgctcgttatccggaccatatgaa<br>gcagcatgacttcttcaagtccgccatgccggaaggctatgtgcaggaacgcacg<br>atttcctttaaggatgacggcacgtacaaaacgcgtgcggaagtgaaatttgaagg<br>cgataccctggtaaaccgcattgagctgaaaggcattgactttaaagaagacggc<br>aatatcctgggccataagctggaatacaattttaacagccacaatgtttacatcacc<br>gccgataaacaaaaaaatggcattaaagcgaattttaaaattcgccacaacgtgg<br>aggatggcagcgtgcagctggctgatcactaccagcaaaacactccaatcggtga<br>tggtcctgttctgctgccagacaatcactatctgagcacgcaaagcgttctgtctaaa<br>gatccgaacgagaaacgcgatcatatggttctgctggagttcgtaaccgcagcgg<br>gcatcacgcatggtatggatgaactgtacaaatgatgataataatctagaccaggc<br>atcaaataaaacgaaaggctcagtcgaaagactgggcctttcgttttatctgttgtttgt<br>cggtgaacgctctctactagagtcacactggctcaccttcgggtgggcctttctgcgtt<br>tataggtacccgtggatcctctagagtcgacctgcaggcatgcaagcttttttacggct<br>agctcagtcctaggtatagtgctagcccagccagagaaacactctttaacagggg<br>gtagtatgatgtctcgtttagataaaagtaaagtgattaacagcgcattagagctgctt<br>aatgaggtcggaatcgaaggtttaacaacccgtaaactcgcccagaagctaggtg<br>tagagcagcctacattgtattggcatgtaaaaaataagcgggctttgctcgacgcctt<br>agccattgagatgttagataggcaccatactcacttttgccctttagaaggggaaag<br>ctggcaagattttttacgtaataacgctaaaagttttagatgtgctttactaagtcatcgc<br>gatggagcaaaagtacatttaggtacacggcctacagaaaaacagtatgaaactc<br>tcgaaaatcaattagcctttttatgccaacaaggtttttcactagagaatgcattatatg<br>cactcagcgcagtggggcattttactttaggttgcgtattggaagatcaagagcatca<br>agtcgctaaagaagaaagggaaacacctactactgatagtatgccgccattattac<br>gacaagctatcgaattatttgatcaccaaggtgcagagccagccttcttattcggcctt<br>gaattgatcatatgcggattagaaaaacaacttaaatgtgaaagtgggtcttaaggc<br>atcaaataaaacgaaaggctcagtcgaaagactgggcctttcgttttatctgttgtttgt<br>cggtgaacgctctcctgagtaggacaaatccgccgccctagacctagggcgttcg<br>gctgcggcgagcggtatcagctcactcaaaggcggtaatacggttatccacagaa<br>tcaggggataacgcatgcaagaacatgtgagcaaaaggccagcaaaaggcca<br>ggaaccgtaaaaaggccgcgttgctggcgtttttccataggctccgcccccctgac<br>gagcatcacaaaaatcgacgctcaagtcagaggtggcgaaacccgacaggact<br>ataaagataccaggcgtttccccctggaagctccctcgtgcgctctcctgttccgacc<br>ctgccgcttaccggatacctgtccgcctttctcccttcgggaagcgtggcgctttctcat<br>agctcacgctgtaggtatctcagttcggtgtaggtcgttcgctccaagctgggctgtgt<br>gcacgaaccccccgttcagcccgaccgctgcgccttatccggtaactatcgtcttga<br>gtccaacccggtaagacacgacttatcgccactggcagcagccactggtaacag<br>gattagcagagcgaggtatgtaggcggtgctacagagttcttgaagtggtggccta<br>actacggctacactagaaggacagtatttggtatctgcgctctgctgaagccagtta<br>ccttcggaaaaagagttggtagctcttgatccggcaaacaaaccaccgctggtagc<br>ggtggtttttttgtttgcaagcagcagattacgcgcagaaaaaaaggatctcaagaa<br>gatcctttgatcttttctacggggtctgacgctcagtggaacgaaaactcacgttaag |

| | | |
|---|---|---|
| | ggattttggtcatgactagtgcttggattctcaccaataaaaaacgcccggcggcaa<br>ccgagcgttctgaacaaatccagatggagttctgaggtcattactggatctatcaac<br>aggagtccaagcgagctcgatatcaaattacgccccgccctgccactcatcgcagt<br>actgttgtaattcattaagcattctgccgacatggaagccatcacaaacggcatgat<br>gaacctgaatcgccagcggcatcagcaccttgtcgccttgcgtataatatttgcccat<br>ggtgaaaacgggggcgaagaagttgtccatattggccacgtttaaatcaaaactg<br>gtgaaactcacccagggattggctgagacgaaaaacatattctcaataaacccttt<br>agggaaataggccaggttttcaccataacacgccacatcttgcgaatatatgtgtag<br>aaactgccggaaatcgtcgtggtattcactccagagcgatgaaaacgtttcagtttg<br>ctcatggaaaacggtgtaacaagggtgaacactatcccatatcaccagctcaccgt<br>ctttcattgccatacgaaattccggatgagcattcatcaggcgggcaagaatgtgaa<br>taaaggccggataaaacttgtgcttatttttctttacggtctttaaaaaggccgtaatatc<br>cagctgaacggtctggttataggtacattgagcaactgactgaaatgcctcaaaatg<br>ttctttacgatgccattgggatatatcaacggtggtatatccagtgatttttttctccattta<br>gcttccttagctcctgaaaatctcgataactcaaaaaatacgaccgg | |
| pKD182 | agcaagcgaaccggaattgccagctggggcgccctctggtaaggttgggaagcc<br>ctgcaaagtaaactggatggctttcttgccgccaaggatctgatggcgcaggggat<br>caagatctgatcaagagacaggatgaggatcgtttcgtatgagggaagcggtgat<br>cgccgaagtatcgactcaactatcagaggtagttggcgtcatcgagcgccatctcg<br>aaccgacgttgctggccgtacatttgtacggctccgcagtggatggcggcctgaag<br>ccacacagtgatattgatttgctggttacggtgaccgtaaggcttgatgaaacaacg<br>cggcgagctttgatcaacgaccttttggaaacttcggcttcccctggagagagcgag<br>attctccgcgctgtagaagtcaccattgttgtgcacgacgacatcattccgtggcgtta<br>tccagctaagcgcgaactgcaatttggagaatggcagcgcaatgacattcttgcag<br>gtatcttcgagccagccacgatcgacattgatctggctatcttgctgacaaaagcaa<br>gagaacatagcgttgccttggtaggtccagcggcggaggaactctttgatccggttc<br>ctgaacaggatctatttgaggcgctaaatgaaaccttaacgctatggaactcgccg<br>cccgactgggctggcgatgagcgaaatgtagtgcttacgttgtcccgcatttggtac<br>agcgcagtaaccggcaaaatcgcgccgaaggatgtcgctgccgactgggcaatg<br>gagcgcctgccggcccagtatcagcccgtcatacttgaagctagacaggcttatctt<br>ggacaagaagaagatcgcttggcctcgcgcgcagatcagttggaagaatttgtcc<br>actacgtgaaaggcgagatcaccaaggtagtcggcaaataagagctcgcttgga<br>ctcctgttgatagatccagtaatgacctcagaactccatctggatttgttcagaacgct<br>cggttgccgccgggcgttttttattggtgagaatccaagcactagtaacaacttatatc<br>gtatggggctgacttcaggtgctacatttgaagagataaattgcactgaaatctagta<br>atattttatctgattaataagatgatcttcttgagatcgttttggtctgcgcgtaatctcttgc<br>tctgaaaacgaaaaaaccgccttgcagggcggtttttcgaaggttctctgagctacc<br>aactctttgaaccgaggtaactggcttggaggagcgcagtcaccaaaacttgtcctt<br>tcagtttagccttaaccggcgcatgacttcaagactaactcctctaaatcaattacca<br>gtggctgctgccagtggtgcttttgcatgtctttccgggttggactcaagacgatagtta<br>ccggataaggcgcagcggtcggactgaacggggggttcgtgcatacagtccagc<br>ttggagcgaactgcctacccggaactgagtgtcaggcgtggaatgagacaaacg<br>cggccataacagcggaatgacaccggtaaaccgaaaggcaggaacaggaga<br>gcgcacgagggagccgccaggggaaacgcctggtatctttatagtcctgtcggg<br>tttcgccaccactgatttgagcgtcagatttcgtgatgcttgtcaggggggcggagcc<br>tatggaaaaacggctttgccgcggccctctcacttccctgttaagtatcttcctggcat<br>cttccgggaaatctccgccccgttcgtaagccatttccgctcgccgcagtcgaacga<br>ccgagcgtagcgagtcagtgagcgaggaagcggaatatatcccgaagcggcat<br>gcatttacgttgacaccatcgaatggtgcaaaacctttcgcggtatggcatgatagc<br>gcccggaagagagtcaattcagggtggtgaatgtgaaaccagtaacgttatacga<br>tgtcgcagagtatgccggtgtctcttatcagaccgtttcccgcgtggtgaaccaggcc<br>agccacgtttctgcgaaaacgcgggaaaaagtggaagcggcgatggcggagct<br>gaattacattcccaaccgcgtggcacaacaactggcgggcaaacagtcgttgctg<br>attggcgttgccacctccagtctggccctgcacgcgccgtcgcaaattgtcgcggcg<br>attaaatctcgcgccgatcaactgggtgccagcgtggtggtgtcgatggtagaacg<br>aagcggcgtcgaagcctgtaaagcggcggtgcacaatcttctcgcgcaacgcgtc | [2] |

| | |
|---|---|
| | agtgggctgatcattaactatccgctggatgaccaggatgccattgctgtggaagct<br>gcctgcactaatgttccggcgttatttcttgatgtctctgaccagacacccatcaacag<br>tattattttctcccatgaagacggtacgcgactgggcgtggagcatctggtcgcattg<br>ggtcaccagcaaatcgcgctgttagcgggcccattaagttctgtctcggcgcgtctg<br>cgtctggctggctggcataaatatctcactcgcaatcaaattcagccgatagcggaa<br>cgggaaggcgactggagtgccatgtccggttttcaacaaaccatgcaaatgctga<br>atgagggcatcgttcccactgcgatgctggttgccaacgatcagatggcgctgggc<br>gcaatgcgcgccattaccgagtccgggctgcgcgttggtgcggatatctcggtagtg<br>ggatacgacgataccgaagacagctcatgttatatcccgccgttaaccaccatcaa<br>acaggattttcgcctgctggggcaaaccagcgtggaccgcttgctgcaactctctca<br>gggccaggcggtgaagggcaatcagctgttgcccgtctcactggtgaaaagaaa<br>aaccaccctggcgcccaatacgcaaaccgcctctccccgcgcgttggccgattcat<br>taatgcagctggcacgacaggtttcccgactggaaagcgggcagtgaggcatca<br>aataaaacgaaaggctcagtcgaaagactgggcctttcgttttatctgttgtttgtcggt<br>gaacgctctcctgagtaggacaaatccgccgccctagacctagggcgttcggctg<br>cggcgagcggtatcagctcactcaaaggcggtaatacgtaaatcactgcataattc<br>gtgtcgctcaaggcgcactcccgttctggataatgtttttttgcgccgacatcataacgg<br>ttctggctaatattctgaaatgagctgttgacaattaatcatcggctcgtataatgtgtg<br>gaattgtgagcggataacaatttcacacaaacgaccatcgacgtccgaccctaac<br>gtatactatgtcccgcctgctgctgtgtatctgtgttctgctgttctcttctgtggcgtggtct<br>aaaccgcagcagttttatgtgggcgtactggctaactggggtcatcagcaagccgtt<br>gaacgttggaccccgatgatggagtatctgaacgaacatgtgccggacgcggaat<br>ttcacgtctacccgggcaacttcaaagcactgaacctggcaatggaactgggcca<br>gattcagttcattatcactaacccgggccaatatctgtacctgagcaatcagtacccg<br>ctgtcttggctggcgaccatgcgttctaagcgtcacgatggtaccacttctgcgatcg<br>gttccgccattattgtccgcgcggacagcgactaccgcaccctgtacgacctgaaa<br>ggtaaagtggtggctgcgtccgacccgcatgctctgggtggctaccaagcgaccgt<br>cggtctgatgcattccctgggcatggatccggacaccttcttcggtgaaaccaagttt<br>ctgggctttccactggatccgctgctgtaccaagttcgtgatggcaacgttgacgcgg<br>ccattaccccactgtgcactctggaggacatggttgcacgcggcgtactgaaatctt<br>ccgattttcgtgtgctgaaccctagccgcccggatggtgtagaatgccagtgctctac<br>caccctgtacccgaactggtctttcgctgcgactgagtctgtatccaccgaactgtcta<br>aagaaatcacgcaggcactgctggaactgccatccgacagcccggcagctatca<br>aagcgcaactgaccggctggaccagcccgatctcccaactggcggtaatcaaac<br>tgttcaaagagctgcacgtaaaaaccccggactctagccgttgggaagccgttaa<br>gaagtggctggaagaaaaccgtcactggggtatcctgtctgttctggtgttcatcattg<br>caacgctgtatcacctgtggattgaataccgcttccaccaaaaaagctcttctctgat<br>cgaatctgaacgtcagctgaaacagcaagctgttgccctggaacgtctgcaatctg<br>ctagcatcgttggtgaaattggtgcgggtctggcccacgagattaatcagccgatcg<br>ctgcaattacctcttattctgaaggtggcatcatgcgcctgcaaggtaaagaacagg<br>cggatacggatagctgcatcgaactgctggaaaaaatccacaaacagagcactc<br>gcgcaggcgaagtggtgcaccgcatccgtggtctgctgaaacgtcgtgaagcggt<br>gatggtagatgttaacatcctgaccctggtggaagaatccatcagcctgctgcgtct<br>ggagctggcacgtcgcgaaatccagatcaacactcagatcaaaggtgaaccgttc<br>ttcattactgccgaccgcgttggcctgctgcaagttctgattaacctgatcaaaaactc<br>cctggacgcgatcgctgaatctgataatgcccgttctggtaaaatcaacatcgaact<br>ggactttaaagagtaccaggtaaacgtctccatcatcgataacggtccgggcctgg<br>cgatggattctgacactctgatggctacgttttacactaccaaaatggatggcctggg<br>cctgggtctggcaatctgccgcgaagttatcagcaaccacgacggccacttcctgct<br>gtccaaccgtgacgacggcgttctgggctgtgtggcaaccctgaatctgaaaaaac<br>gcggttctgaagtgccgatcgaagtctaataatctagaccaggcatcaaataaaac<br>gaaaggctcagtcgaaagactgggcctttcgttttatctgttgtttgtcggtgaacgctc<br>tctactagagtcacactggctcaccttcgggtgggcctttctgcgtttataggtacccg<br>gggatcctctagagtcgacctgcaggcatgcaagctt |

**Table S4.** Strains used in this study.

| # | Title | Description | Cell type | Antibiotics |
|---|---|---|---|---|
| 1 | 182+184 | PKD182 + PKD184 For positive thiosulfate controls | E. coli NEB5a | Cm/Sm |
| 2 | 182 + 189 | PKD182 + PKD189 for positive thiosulfate controls | E. coli NEB5a | Cm/Sm |
| 3 | Mut1-1 + 184 | Mut1-1 + PKD184 biosensor (from round 1 of directed evolution) | E. coli NEB5a | Cm/Sm |
| 4 | Mut1-2 + 184 | Mut2-1 + PKD184 biosensor (from round 1 of directed evolution) | E. coli NEB5a | Cm/Sm |
| 5 | PC1+184 | CCNY PC1 + pkd184 (from initial computational design) | E. coli NEB5a | Cm/Sm |
| 6 | PC2+184 | CCNY PC2 + pkd184 (from initial computational design) | E. coli NEB5a | Cm/Sm |
| 7 | PC3+184 | CCNY PC3 + pkd184 (from initial computational design) | E. coli NEB5a | Cm/Sm |
| 8 | PC4+184 | CCNY PC4 + pkd184 (from initial computational design) | E. coli NEB5a | Cm/Sm |
| 9 | PC5+184 | CCNY PC5 + pkd184 (from initial computational design) | E. coli NEB5a | Cm/Sm |
| 10 | PC6+184 | CCNY PC6 + pkd184 (from initial computational design) | E. coli NEB5a | Cm/Sm |
| 11 | PC7+184 | CCNY PC7 + pkd184 (from initial computational design) | E. coli NEB5a | Cm/Sm |
| 12 | PC8+184 | CCNY PC8 + pkd184 (from initial computational design) | E. coli NEB5a | Cm/Sm |
| 13 | PC9+184 | CCNY PC9 + pkd184 (from initial computational design) | E. coli NEB5a | Cm/Sm |
| 14 | PC10+184 | CCNY PC10 + pkd184 (from initial computational design) | E. coli NEB5a | Cm/Sm |
| 15 | Mut2-1 + 184 | Mut2-1 + PKD184 biosensor (from round 2 of directed evolution) | E. coli NEB5a | Cm/Sm |
| 16 | Mut2-2+ 184 | Mut2-2 + PKD184 biosensor (from round 2 of directed evolution) | E. coli NEB5a | Cm/Sm |
| 17 | Mut2-3 + 184 | Mut2-3 + PKD184 biosensor (from round 2 of directed evolution) | E. coli NEB5a | Cm/Sm |
| 18 | Mut2-4 + 184 | Mut2-4 + PKD184 biosensor (from round 2 of directed evolution) | E. coli NEB5a | Cm/Sm |
| 19 | Mut3-1 + 184 | Mut3-1 + PKD184 biosensor (from round 3 of directed evolution) | E. coli NEB5a | Cm/Sm |
| 20 | Mut3-2 + 184 | Mut3-2 + PKD184 biosensor (from round 4 of directed evolution) | E. coli NEB5a | Cm/Sm |
| 21 | Mut3-3 + 184 | Mut3-3 + PKD184 biosensor (from round 5 of directed evolution) | E. coli NEB5a | Cm/Sm |

| | | | | |
|---|---|---|---|---|
| 22 | Mut3-4 + 184 | Mut3-4 + PKD184 biosensor (from round 6 of directed evolution) | E. coli NEB5a | Cm/Sm |
